# Sequences and proteins that influence mRNA processing in *Trypanosoma brucei*: evolutionary conservation of SR-domain and PTB protein functions

**DOI:** 10.1101/2022.04.25.489340

**Authors:** Albina Waithaka, Olena Maiakovska, Dirk Grimm, Larissa Melo do Nascimento, Christine Clayton

**Affiliations:** Heidelberg University Centre for Molecular Biology (ZMBH), Im Neuenheimer Feld 282, D-69120 Heidelberg, Germany; Department of Infectious Diseases/Virology, Medical Faculty, University of Heidelberg, BioQuant, Im Neuenheimer Feld 267, D-69120 Heidelberg, Germany; German Center for Infection Research (DZIF) and German Center for Cardiovascular Research (DZHK), partner site Heidelberg, D-69120 Heidelberg, Germany

**Keywords:** *Trypanosoma brucei*, *trans* splicing, polyadenylation, polypyrimidine tract, Polypyrimidine tract binding protein, SR-domain protein.

## Abstract

Spliced leader *trans* splicing is the addition of a short, capped sequence to the 5’ end of mRNAs. It is widespread in eukaryotic evolution, but factors that influence *trans* splicing acceptor site choice have been little investigated. In Kinetoplastids, all protein-coding mRNAs are 5’ *trans* spliced. A polypyrimidine tract is usually found upstream of the AG splice acceptor, but there is no branch point consensus; moreover, splicing dictates polyadenylation of the preceding mRNA. We here describe a *trans* splicing reporter system that can be used for studies and screens concerning the roles of sequences and proteins in processing site choice and efficiency. Splicing was poor with poly(U) tracts less than 9 nt long, and was influenced by nearby secondary structures. A screen for signals resulted in selection of sequences that were on average 45% U and 35% C. Tethering of either the splicing factor SF1, or the cleavage and polyadenylation factor CPSF3 within the intron stimulated processing in the correct positions, while tethering of two possible homologues of Opisthokont PTB inhibited processing. In contrast, tethering of SR-domain proteins RBSR1, RBSR2, or TSR1 or its interaction partner TSR1IP, promoted use of alternative signals upstream of the tethering sites. RBSR1 interacts predominantly with proteins implicated in splicing, whereas the interactome of RBSR2 is more diverse. These results suggest that the functions of PTB and SR-domain proteins in splice site definition were already present in the last eukaryotic common ancestor.

## Introduction

Processing of nuclear-encoded mRNAs by 5’ *trans* splicing of short capped “spliced leader” sequences is scattered throughout eukaryotic evolution, being found in at least three different supergroups (Krchňáková et al. 2017). The mechanism is overall similar to that of *cis* splicing; and while it is usually assumed to have evolved independently numerous times, there is also a possibility that it was present in the common ancestor and subsequently lost (Krchňáková et al. 2017). In comparison with *cis* splicing, both the sequences required for *trans* splicing, and the factors that regulate it have been relatively little investigated.

In *Trypanosoma brucei* and related kinetoplastid parasites, dependence on *trans* splicing is extreme, since transcription of nearly all protein-coding genes is polycistronic. Trypanosomes therefore serve as a useful model system to study *trans* splicing. The mature capped 5’-ends are formed by addition of the capped spliced leader (*SL*) (reviewed in (Michaeli 2011; Preusser et al. 2012). The spliced leader precursor mRNAs, which are called *SLRNA*s, include the *SL* sequence and a short intron. The *SLRNA* assembles into a ribonucleoprotein particle (RNP) (Cross et al. 1991) which is recruited with other spliceosomal components to the splice acceptor sites of pre-mRNAs (Figure 1A). As in other eukaryotes, splicing factors are thought to recognize a polypyrimidine tract (PPT) upstream of the splice site. The spliced leader intron forms a Y structure by 2’-5’ linkage with any A residue immediately upstream of the PPT; there is no branch-point consensus sequence (Lücke et al. 2005). Polyadenylation occurs approximately 100 nt upstream of the PPT. There is no polyadenylation signal within the 3’-untranslated region (UTR) but A residues are preferred at the cleavage site (Hug et al. 1994; Matthews et al. 1994; Schürch et al. 1994; Vassella et al. 1994). Subsequently the Y is debranched and the intergenic sequence is discarded. Splicing and polyadenylation are physically and mechanistically linked: if splicing is prevented, polyadenylation also cannot occur, and *vice versa* (Ullu et al. 1993; Hendriks et al. 2003; Clayton and Michaeli 2011; Begolo et al. 2018; Wall et al. 2018). The components of the splicing and polyadenylation complexes have been characterized (Palfi et al. 1991; Cross et al. 1993; Lucke et al. 1997; Palfi et al. 2000; Palfi et al. 2002; Preusser et al. 2009; Jae et al. 2010; Koch et al. 2016) but the nature of the connection between them is still not understood.

**Figure 1.**
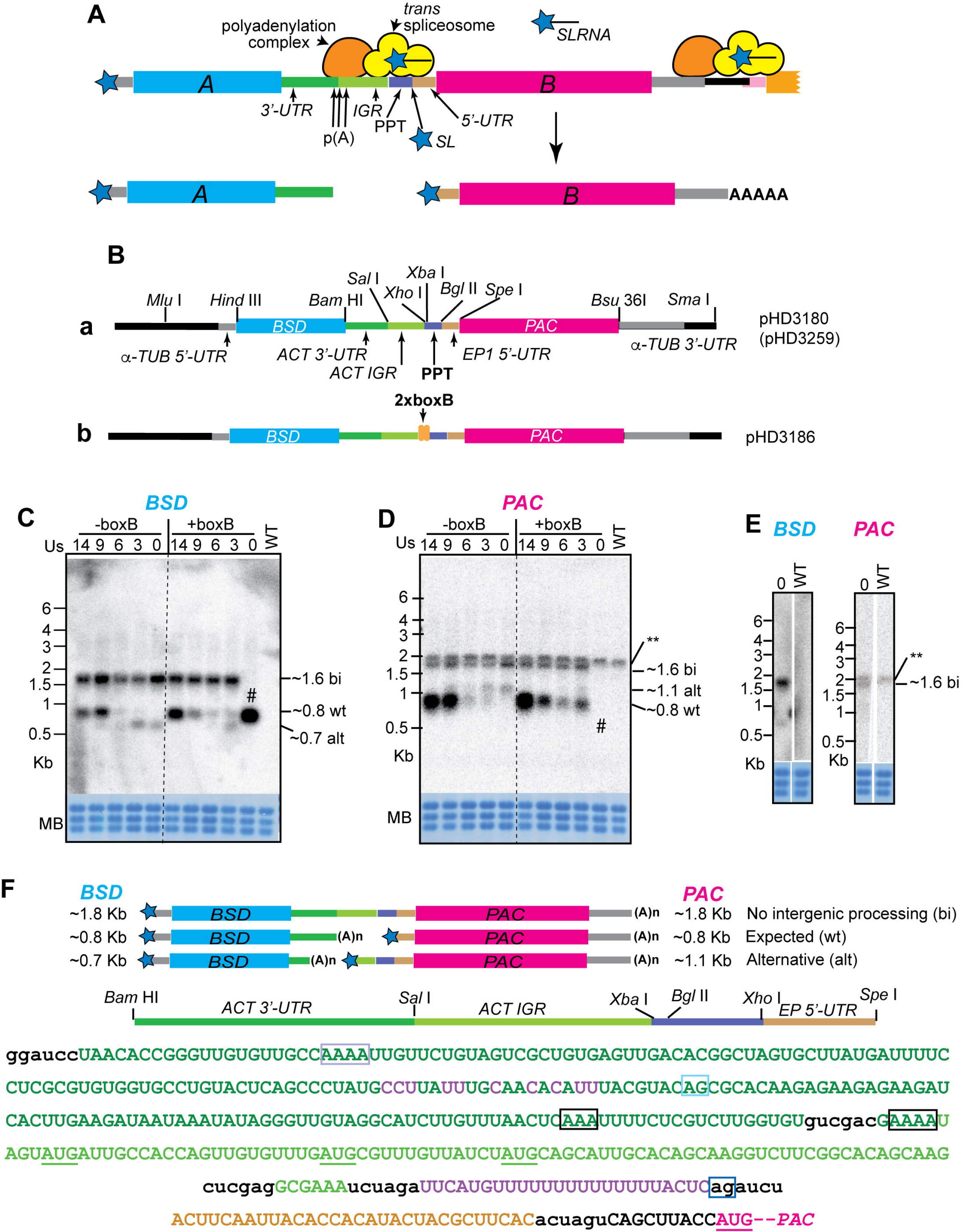
Effect of PPT length and a secondary structure on mRNA processing. A. Scheme showing the mechanism of *trans* splicing and polyadenylation. Important players and sequence regions are labelled. B. Maps of the two vectors used including important restriction sites. C. Northern blot showing examples of *BSD* mRNAs using different PPT lengths. All results (except those for +BoxB with no PPT) were confirmed using at least three independent transformed populations. The methylene blue-stained rRNA shows the loading. WT are cells that do not have the plasmid. D. Northern blot for *PAC* mRNA. The WT lane shows a band labelled ** that cross-hybridised with the *PAC* probe. This appeared with varying intensities. E. *BSD* and *PAC* mRNAs from a +BoxB, 0-PPT clone which retained the *PAC* gene. F. Sequence between the *BSD* and *PAC* genes. The upper maps show the three different processing patterns: the partially processed bicistronic mRNA with no cleavage between the two open reading frames; the expected processing pattern; and the alternative processing pattern. The map of the intergenic region is expanded between the mRNA maps and the sequence. The sequence is colour-coded like the maps, and restriction sites are in lower case. The processing sites are surrounded by boxes (black for expected poly(A) site, dark blue for expected splice acceptor, grey for alternative poly(A), cyan for alternative splice acceptor). Start codons between the alternative and the wild-type PPT are underlined in green, and the wild-type *PAC* start codon in magenta.

The sequence requirements for kinetoplastid *trans* splicing have been investigated in various ways. Many of the initial experiments were done by transient transfection of reporters, followed by poly(A) and splice site mapping by Northern blotting or reverse-transcription followed by PCR (Hug et al. 1994; Matthews et al. 1994; Schürch et al. 1994; Vassella et al. 1994). These experiments revealed that if the PPT that normally defined the splice site had been removed, an alternative sequence was used instead. Studies of splicing and polyadenylation sites by RNA sequencing (RNASeq) showed that many mRNAs use more than one AG splice acceptor site (Kolev et al. 2010). Two life-cycle stages of *T. brucei* grow well *in vitro*: the mammalian “bloodstream” form, and the “procyclic” form which grows in the midgut of Tsetse flies. One of the early RNASeq studies revealed developmentally regulated splice sites (Nilsson et al. 2010). Analysis of the RNASeq results also revealed that the lengths of PPTs vary from 9-40 nt; most are between 12 and 24 nt with a predominance of U over C. The distances between the preceding poly(A) sites and the PPT are usually 100-150 nt, and use of several different poly(A) sites is the norm. Siegel *et al*. (Siegel et al. 2005) used a transiently transfected reporter, with RNA polymerase I transcription, to study the effects of PPT length and composition on *trans* splicing. They inserted various 17mers at the position of the PPT and then measured protein production from the resulting *trans* spliced RNA. Maximum efficiency was obtained with (U)_20_. (C)_17_ was inactive, but mixtures of Cs and Us were as good as Us alone. Insertions of purines, however, had clear deleterious effects.

Trypanosome precursor mRNAs, especially 3’-untranslated regions, contain numerous polypyrimidine tracts that are not normally used for *trans* splicing. Moreover, various studies have suggested that trypanosome mRNAs show differences in processing efficiency. Modelling of mRNA levels, based on high-throughput measurements, suggested that mRNA turnover rates and gene copy numbers could not account for all variations in mRNA abundance (Fadda et al. 2014; Antwi et al. 2016). The authors suggested that if an mRNA precursor is inefficiently processed, some of the precursors will be destroyed in the nucleus by quality control mechanisms, resulting in low mRNA abundance. In Opisthokonts, splice acceptor site choice is governed, at least in part, by specific RNA-binding proteins: Polypyrimidine tract binding proteins (PTBs) inhibit splicing (Keppetipola et al. 2012), whereas some proteins with serine-arginine-rich (SR) domains define exons through binding downstream of splice acceptor sites (Howard and Sanford 2015). Results from studies of selected trypanosome RNA-binding proteins and basal splicing factors indeed indicated that different factors control specific mRNAs. Possible regulators include DRBD3 and DRBD4, possible homologues of PTBs (Estévez 2008; Stern et al. 2009); an HNRNPF/H homologue (Gupta et al. 2013); and various SR-domain proteins including TSR1 and its binding partner TSR1IP (Ismaili et al. 1999; Ismaili et al. 2000; Gupta et al. 2014). When expression of any of these proteins was reduced, some mRNAs increased and others decreased. The results suggested that depletion of the proteins affected the stabilities of some mRNAs, as well as processing of others. However, detailed effects on mRNA processing were not actually demonstrated. The different effects presumably depend at least partially on whether the protein binds in the intron or in the body of the mature mRNA (Gupta et al. 2013; Das et al. 2015) but no binding sites in precursors have been identified.

In this paper, we describe a screen for active PPTs, and demonstrate the effects of various splicing factors or regulators when tethered immediately upstream of the PPT. The results give useful insights into both the sequences required for active *trans* splice acceptor sites, and the functions of SR-domain proteins and PTB homologues in determining splice site choice.

## Materials and Methods

### Trypanosome culture and modification

The experiments in this study were carried out using monomorphic *T. brucei* Lister 427 bloodstream form parasites constitutively expressing the Tet repressor (Alibu et al. 2005). The parasites were routinely cultured at 37°C in HMI-9 medium supplemented with 10% heat-inactivated fetal bovine serum (v/v), 1% (v/v) penicillin/streptomycin solution (Labochem international, Germany), 15 µM L-cysteine, and 0.2 mM β-mercaptoethanol in the presence of 5% CO_2_ and 95% humidity. During proliferation, the cells were diluted to 1×10^5^ cells/ml and maintained between 0.2-2×10^6^ /ml. Cell densities were determined using a Neubauer chamber. For generation of stable cell lines, ∼1-2×10^7^ cells were transfected by electroporation with 10 µg of linearized plasmid at 1.5 kV on an AMAXA Nucleofector. Selection of new transfectants was done after addition of 5 μg/ml blasticidin (InvivoGen). Independent populations were obtained by serial dilution about 6 h after transfection. RNAi or tagged protein expression was induced using 100 ng/ml tetracycline. The primers and plasmids used are listed in Supplementary Table S1. Selected sequences are found in Supplementary Text 1 and the reporter plasmids are available from the European Plasmid repository (https://www.plasmids.eu).

### PPT library preparation

Starting with pHD3180, 11 random nucleotides were inserted in place of the 14 Ts at the “PPT” position, by site directed mutagenesis using the primers in Supplementary Table S1. The resulting pool of plasmids was desalted by ethanol precipitation and transformed into TOP10 One Shot electrocompetent *E. coli* cells (ThermoFisher) by electroporation using 0.2 cm pre-chilled cuvettes. The electroporation was done at, 2.5 kV, 25 μF, 200 Ω TC +/-4.5 milliseconds. After transformation, the bacteria were plated on LB plates and incubated for 16-17 h at 37°C, after which the colonies were picked and expanded overnight in 25 ml LB media and plasmids extracted. The plasmid library was transfected into *T. brucei* and selected with blasticidin as above. Surviving clones were then grown in 1x (200 μg/ml) puromycin. Genomic DNA from the original population and the puromycin-selected μ parasites was extracted and the region targeting PPT amplified by PCR. The amplicons were run on a 3% agarose gel, and the products were cut, purified, and sent for sequencing (see below).

### RNA manipulation

RNA was prepared using Trizol according to the manufacturer’s instructions. For Northern blots, 10 µg of total RNA was loaded per lane on formaldehyde denaturing gels, and blotted onto Nytran+. The membrane was stained with methylene blue and scanned before hybridization with [^32^P]-labelled probes. The signals were detected by phosphorimaging.

### DNA sequencing and data analysis

For DNA and RNA sequencing, libraries were prepared by David Ibberson at the CellNetworks Deep Sequencing Core Facility at the University of Heidelberg, followed by sequencing at EMBL.

For the PPT analysis, the relevant integrated plasmid sequence was amplified (Supplementary Table S1) from total population DNA and then sequenced (Supplementary Table S1). Raw paired Illumina (Miseq) reads were merged using the fastq-join tool with default parameters (version 1.3.1). A custom bash code was used to subset PPT sequences by searching for the flanking sequences (GGCGAAATCTAGA and AGATCTACTTC). The number of unique PPT sequences, their lengths and composition were counted and used for further visualization in R (version 3.6.3; R Core Team, 2022) on Ubuntu 20.04.2 LTS. The dplyr (version 1.0.8) (Wickham et al. 2022), tidyverse (Wickham et al. 2019) and ggplot2 (Wickham 2016) packages were applied in data manipulation and plotting in R.

Total RNA was either poly(A)-selected using the standard Illumina kit, or subjected to rRNA depletion using oligonucleotides and RNase H (Minia et al. 2016). RNASeq data were aligned to the TREU927 and Lister 427 (2018) genomes using a custom pipeline that includes Bowtie2 (Langmead and Salzberg 2012; Leiss et al. 2016) and then statistically analysed using a DeSeq2-based pipeline (Love et al. 2014; Leiss and Clayton 2016). Subsequent analyses were done mainly in Microsoft Excel.

### Drug sensitivity tests

To obtain the EC_50_s of puromycin, the drug was serially diluted in 100 µl culture medium in 96 well opaque plates to give concentrations between 50 ng/ml and 26 mg/ml. Trypanosomes were added at a final density of 2×10^4^/ml (100 µl) and incubated for 48 h. After incubation, 20 µl of 0.49 mM Resazurin sodium salt in PBS was added to each well and plates were incubated for a further 24 h. Plates were read on a BMG FLUOstar OPTIMA microplate reader (BMG Labtech GmbH, Germany) with λ_excitation_ = 544 nm and λ_emission_ = 590 nm, to assess the number of surviving viable cells (Raz et al. 1997; Sykes and Avery 2009). GraphPad Prism 6 was used to calculate EC_50_. The percentage growth inhibition was plotted against the log_10_ puromycin concentration and analysed in sigmoidal dose response, variable slope mode.

### Mass spectrometry

Proteins associated with RBSR1 and RBSR2 were identified exactly as described in (Falk et al. 2021). Briefly, RBSR1-TAP, RBSR2-TAP and TAP-GFP were purified from cell extracts over IgG beads, then released using His-tagged TEV protease. The latter was subsequently removed using a nickel column. The purified proteins were then run on a polyacrylamide gel and subjected to mass spectrometry. Data were analyzed using PERSEUS (Tyanova et al. 2016).

## Results

### Construction of the reporter

The reporter used for our studies is shown in Figure 1B(a). After restriction site cleavage, it integrates into the tubulin gene repeat, replacing an alpha-tubulin open reading frame. Transcription will therefore be effected by RNA polymerase II. The first open reading frame, encoding blasticidin S deaminase (BSD), is preceded by the native alpha-tubulin 5’-UTR and upstream splicing signals. After the *BSD* gene comes a truncated version of the 3’-UTR from the actin-B (*ACT*) gene, followed by the downstream intergenic region (IGR). The polyadenylation site is fortuitously marked by a *Sal* I site. The PPT from the actin intergenic region is separated from the IGR by two unique restriction sites, and is followed by a unique *Bgl* II site. Finally, a fragment of the *EP* procyclin 5’-UTR, with a few additional nucleotides, precede an open reading frame encoding puromycin acetyltransferase (PAC), and the alpha-tubulin 3’-UTR. Two versions of this plasmid are available, pHD3180 and pHD3259: they differ only in that pHD3259 has a shorter segment of the alpha-tubulin locus upstream of *BSD*. After transcription of this plasmid, the *BSD* mRNA has an alpha-tubulin 5’-UTR and a truncated *ACT* 3’-UTR, while the *PAC* mRNA has the *EP1* 5’-UTR (plus additional sequence) and the alpha-tubulin 3’-UTR. Reconstructed sequences, verified in all critical regions, are available in the supplement.

### The effect of changing the PPT on mRNA processing

First, we examined the effect of changing the PPT. Results are shown in Figure 1C. It was immediately apparent that processing between the *PAC* and *BSD* mRNAs was somewhat inefficient: an mRNA that hybridised with both open reading frame probes (labelled “bi” in all subsequent Figures) was always present. We assume that this RNA is *trans* spliced upstream of *BSD* and polyadenylated downstream of *PAC*, but we have not verified this experimentally. We also do not know whether the bicistronic mRNA is exported to the cytoplasm. Similar bicistronic tubulin mRNAs, which also result from incomplete processing, are usually substrates for degradation by the RNA exosome, most of which is in the nucleus (Kramer et al. 2016). RNAs containing two or more open reading frames also accumulate after inhibition of splicing or polyadenylation (Muhich and Boothroyd 1988; Ullu et al. 1993; Hendriks et al. 2003), and can be exported, at least partially (Kramer et al. 2012).

A precursor with 14 Us corresponds to the sequence originally published for the actin locus (Ben Amar et al. 1988). This yielded the bicistron and processed mRNAs of approximately the expected sizes. Both mRNAs are expected to be about 850 nt long, assuming use of the endogenous tubulin signals (Kimmel et al. 1985) and a mean poly(A) tail of 60 nt (Figure 1C, D). (U)_9_ gave a similar result, but with (U)_6_ and (U)_3_ an mRNA that was about 100 nt shorter was seen. Reducing the PPT length to 9 nt made no difference to the splicing pattern, but further shortening caused the appearance of a weak *PAC* band migrating at about 1.1 kb (Figure 1C, D).

To find out precisely how the PPT changes were affecting processing, we amplified the *PAC* and *BSD* mRNAs and sequenced the products. Results are illustrated in Figure 1F. The top map shows the bicistronic mRNA, which has been processed using only the tubulin signals, and the next map down shows the products from the original construct, with (U)_14_. Sequencing of these products revealed that they had indeed been processed at exactly the expected positions (black box in the sequence for poly(A), dark blue for the splice acceptor). With the shortened PPTs, alternative processing occurred. Now, the *BSD* poly(A) site was 20 nt downstream of the *BSD* cassette (lilac box), giving an mRNA that was predicted to be 200 nt shorter than the original. The alternative *PAC* mRNA splice site was correspondingly 104 nt downstream of the new poly(A) site, *i.e.*, 200 nt upstream of the original one. The longest PPT upstream of the new splice site is UUUUCCUC but this is only 43 nt downstream of the new poly(A) site, which would be unusually close. Other pyrimidines in more appropriate positions are highlighted in purple. The new *PAC* mRNA has an extended 5’-UTR which contains three initiation codons (underlined), with downstream open reading frames of 8, 18 and 23 codons and terminating upstream of the *PAC* initiation codon. We would expect the presence of these “upstream open reading frames” to impair translation of the *PAC* gene severely.

We next looked at the effects of PPT changes on puromycin resistance, measuring survival after two days of treatment using a live-cell fluorescence assay. A typical set of curves is shown in Figure 2A and results for three replicates are in Figure 2C. As expected, puromycin resistance depended on production of mRNA with the short 5’-UTR. Decreasing the length of the PPT from 14 to 9 nt had no effect on puromycin resistance and with no PPT the EC_50_ was two orders of magnitude lower. Cells with (U)_6_ were marginally more resistant than those with (U)_3_, but still with a ten times lower EC_50_ than the (U)_14_ control. These results so far showed that the vector can be used not only to test the effects of specific intergenic regions, PPTs, or 5’-UTRs on gene expression, but also for untargeted screens.

**Figure 2.**
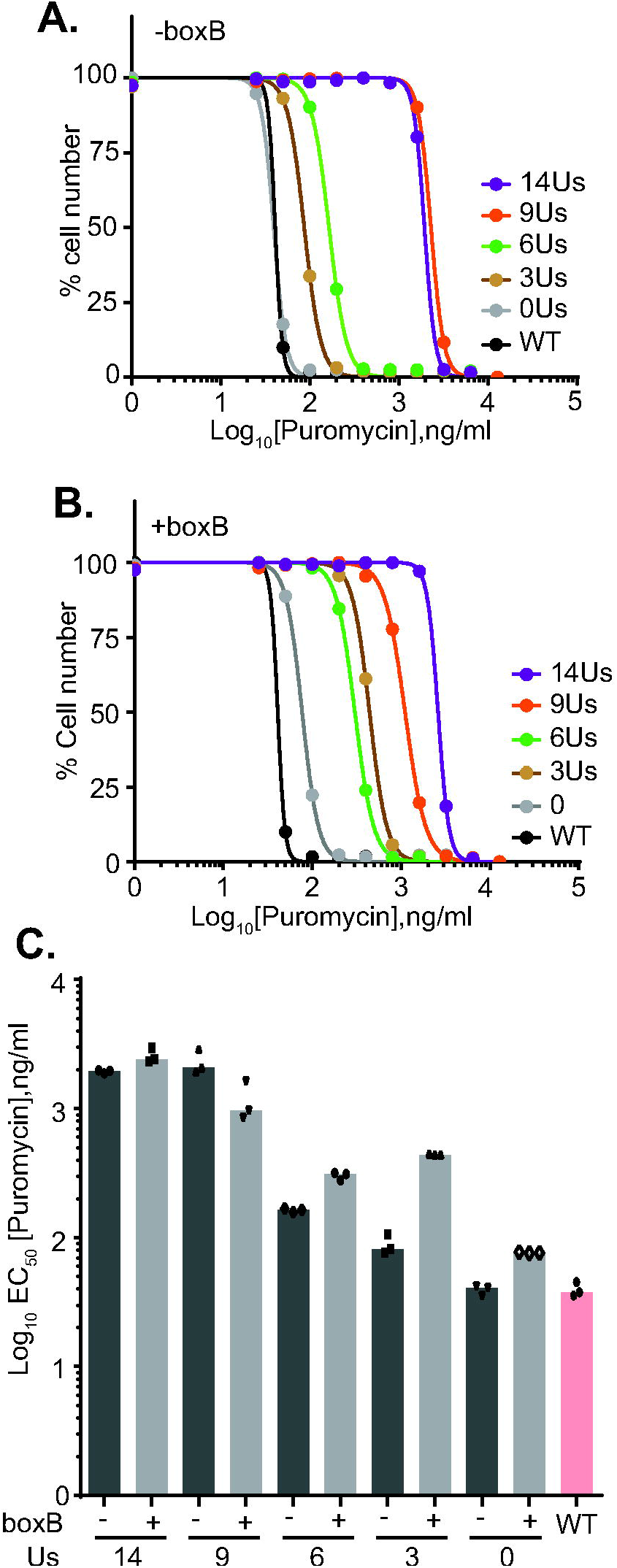
Effect of different PPT lengths on puromycin sensitivity. A. Puromycin sensitivity curves for trypanosomes containing pHD3180 (no boxB) with different PPT lengths. Numbers of viable cells remaining after 48 h (as measured using rezazurin) relative to untreated cells are shown on the y-axis, and the puromycin concentration on the x-axis. B. As (A) but for trypanosomes containing pHD3186 (with boxB) with different PPT lengths. C. Quantitation of three technical replicates for clones shown in Figure 1. Additional clones gave similar results.

### Addition of a boxB sequence upstream of the PPT promotes splicing

The boxB sequence is a 19 nt stem-loop RNA from bacteriophage lambda. It binds to a 22-residue peptide from the lambdaN protein with high affinity. The boxb-lambdaN combination that can be used to “tether” any protein of interest artificially to an RNA (Baron-Benhamou et al. 2004). Since we wished to investigate the effects of potential splicing regulators on our reporter, we incorporated two copies of boxB immediately upstream of the PPT. We then tested the effects on mRNA processing (Figure 1C-E) and puromycin resistance (Figure 2B, C). The presence of 2xboxB had no effect for (U)_14_ but slightly inhibited processing with (U)_9_. Interestingly, insertion of 2xboxB upstream of (U)_3_ yielded results that were similar to those from (U)_6_.

With no PPT at all, Northern blots combined with PCR using genomic DNA showed that four of five clones tested had deleted the *PAC* gene entirely, giving just the normal *BSD* transcript (# in Figure 1C and D). Only one clone had retained the *PAC* gene, and had only the bicistronic *BSD-PAC* mRNA (Figure 1E). This suggests that 2xboxB requires the 3 Us in order to give “correct’ splicing, and has no activity by itself. For quantification of puromycin resistance, this clone was used. The *PAC* gene deletion also occurred with two of the five clones with no PPT and without boxB, and in one clone with (U)_3_ and no boxB. Since all clones were selected with blasticidin, this suggests that the bicistronic RNA alone gives very poor blasticidin resistance, consistent with either retention in the nucleus, or repression of translation caused by the *PAC* sequence in the 3’-UTR.

### Different 5’ regions had small effects on mRNA yields

We next studied the functions of regions upstream of five different genes. In each case, we amplified the region from the polyadenylation site to the nucleotide immediately before the start codon, including the intergenic region with splicing signals and the 5’-UTR (Supplementary Text 1). We used the fragments to replace the region between the *Sal* I and *Spe* I sites in our starting plasmid (Figure 3A).

**Figure 3.**
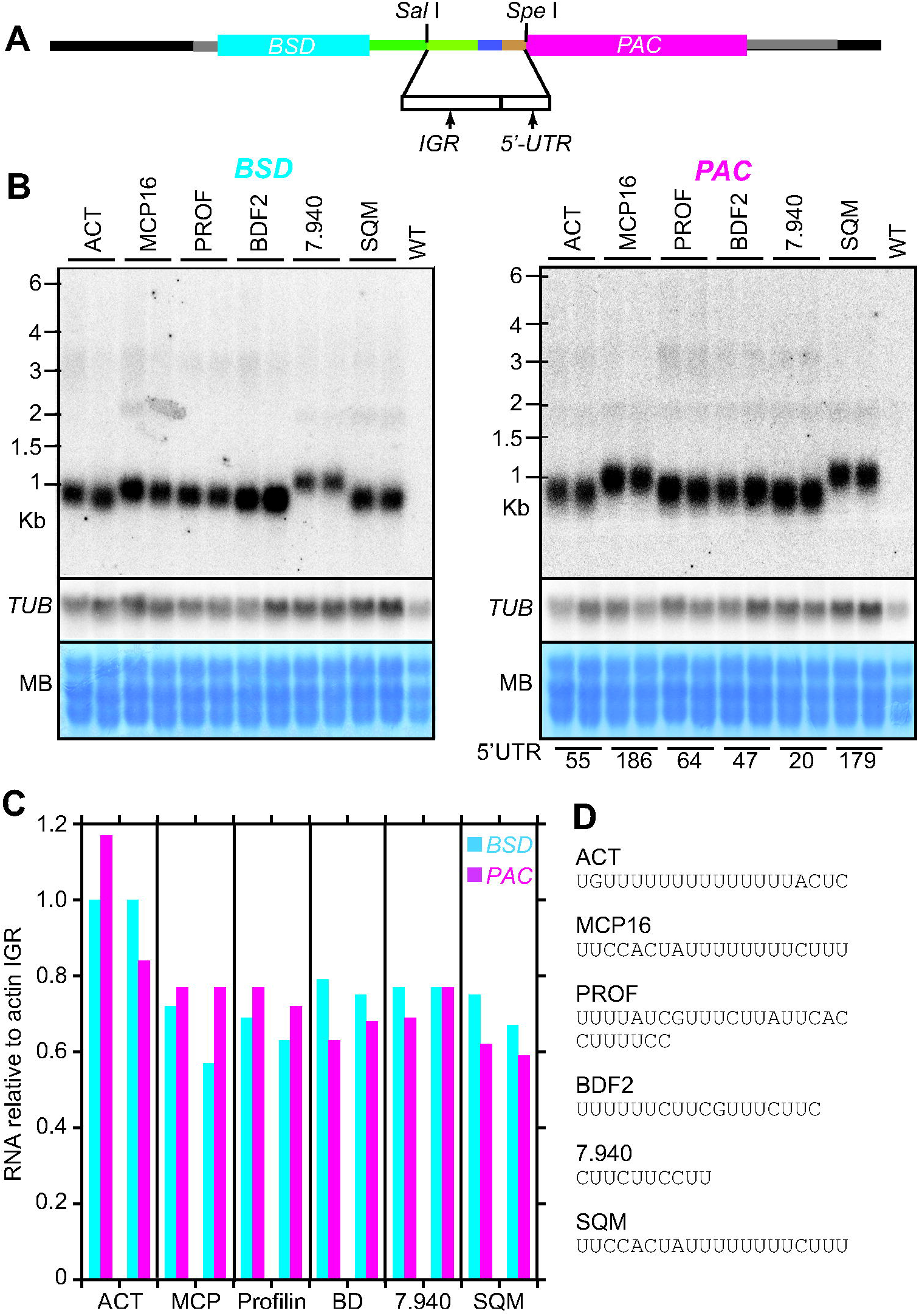
Different intergenic regions and 5’-UTRs. A. Cloning scheme showing positions of the tested sequences. B. Northern blots as in Figures 1C and 1D. The blot was hybridised with a beta-tubulin probe for normalization. Results of two different populations were used. The predicted 5’-UTR lengths are noted below the lanes. C. Amounts of mRNA, obtained by scanning the Northern blot images. Amounts were calculated for two different Northern blots for each population and normalised to the tubulin signal. D. Polypyrimidine tracts upstream of the splice sites. The full sequences are shown in Supplementary Text 1.

As a control, we used the region upstream of the actin gene *ACTA*. A single *PAC* mRNA of the expected size was obtained, while the *BSD* mRNA was also the same size as before, again as expected (Figure 3B). Processing was notably more efficient than it had been for the starting plasmid, since no bicistronic mRNAs were visible. This suggests that the patchwork of different sequences and/or the added restriction sites in pHD3180 had impaired processing efficiency.

The other genes were chosen because their measured mRNA abundances differed in various ways from those that were predicted on the basis of their half-lives. Tb927.7.3940 (MCP16) encodes a mitochondrial carrier protein and was predicted to have an average *trans* splicing efficiency. Tb927.11.13780 (PROF), encoding profilin, and Tb927.7.940, encoding a protein of unknown function, were predicted to be spliced faster than average. In contrast, Tb927.10.7420, encoding bromodomain factor 2 (BDF2), and Tb927.11.3270 (Squaline monooxygenase, SQM) were predicted to be processed slower than average. The mRNA sizes were approximately as predicted (Figure 3B). In one case (Tb927.7.940), the *BSD* mRNA extended beyond the *Sal* I site junction because our cloned fragment turned out to contain about 140 nt of the preceding 3’-UTR. Changing the intergenic region had very little effect on the abundances of the *PAC* or *BSD* mRNAs: all of the different upstream regions gave 20-40% less mRNA than the actin controls (Figure 3C). Thus, if these mRNAs really are spliced with different efficiencies, the sequences we used do not contain the requisite information. One possibility is that the beginning of the coding region might also play a role. Alternatively, the high-throughput mRNA half-life measurements that were used to predict the abundances of the mRNAs may have been incorrect.

### A screen for splicing signals

Next, we made a library of plasmids containing different 11mers in the PPT position. The RNA sequence that results is UCUAGANNNNNNNNNNNAAGAUCU, so that the first available splice acceptor site is immediately downstream of the 11mer. A second one is located 37 nt downstream, 10 nt upstream of the AUG. The plasmids were transfected into trypanosomes and selected with blasticidin. The resulting population was grown in puromycin. DNA was prepared, and the PPT regions from different populations were amplified using upstream and downstream primers (Supplementary Table S1) and then sequenced. An initial trial revealed selection of specific sequences at 0.2 µg/ml or 1 µg/ml puromycin (Supplementary Table S2), and no survival at 2 µg/ml. We therefore repeated the experiment with three independent replicates, first with 0.2 µg/ml puromycin (the normal concentration, 1x in Figure 4), which initially inhibited growth before recovery of the population, and then 1 µg/ml drug (5x in Figure 4), which did not detectably retard replication. Sequencing of the amplicons yielded 2.5-3.3 million reads per population, with up to 20,731 unique sequences in a replicate and coverage from 1x to 790,634x. Selection for 11mers only revealed 30,229 different 11mers, of which 28,248 were present as less than ten reads in the initial population, and 11,135 were completely absent. To find sequences that were lost during selection, we concentrated on those that were initially present as at least ten reads, and for which the median normalized values after puromycin selection were at least five-fold lower. This yielded 1,177 11mers (Figure 4A) with very little base preference at any position, apart from a very weak preference for G and A towards the 3’-end (Figure 4B). Intriguingly, however, 120 of them contained a PPT of at least 6 nt. Separate examination of this subset revealed that the PPTs were concentrated towards the 5’-end of the 11mer (Figure 4B).

**Figure 4.**
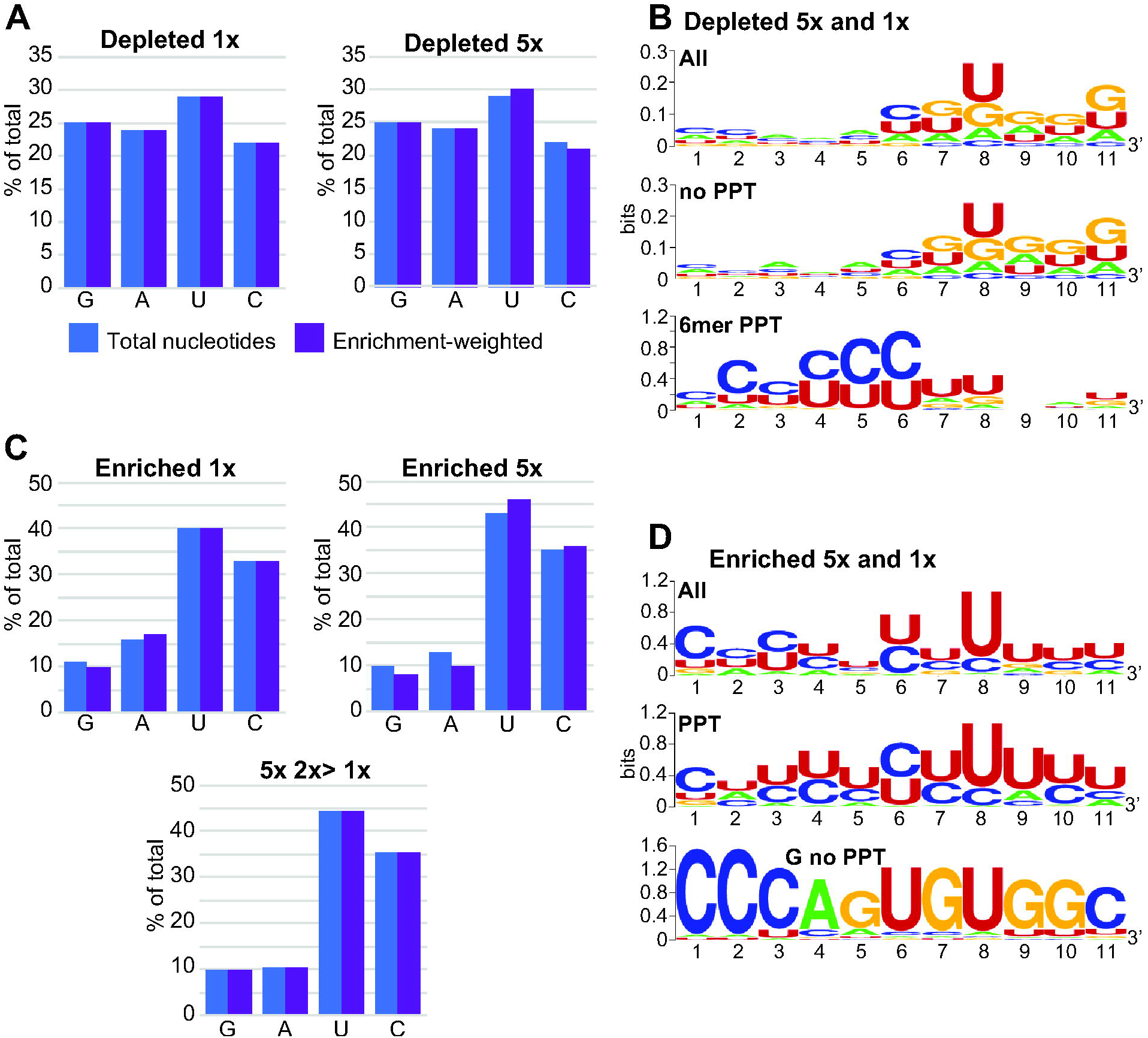
11mer sequences that affect reporter expression. Trypanosomes carrying the reporter (no boxB) with random 11mers in the PPT position were first selected with blasticidin and then with puromycin. Inserts of the starting and selected populations were amplified and sequenced. A. Nucleotide compositions of depleted sequences. We selected sequences that were present at least ten times in the starting population. To determine enrichment in the starting population, the normalized number of reads (reads per million reads, RPM) was then divided by the (median RPM +1) for three populations selected with 0.2 µg/ml or 1 µg/ml puromycin (Depleted 1x and Depleted 5x, respectively). Sequences for which the enrichment in the starting population was at least five-fold were selected. The blue bars show the base compositions if every selected sequence was counted once. The purple bars show the base compositions after each sequence was weighted according to the degree of selection (multiplied by the enrichment factor). B. Sequences that were depleted after both 1x and 5x selection were analyzed using WebLogo (Crooks et al. 2004). The top panel shows all sequences, the middle panel those that lack six contiguous pyrimidines, and the bottom panel those that had six contiguous pyrimidines. C. Base compositions of sequences that were enriched at least five-fold after growth in 1x or 5x puromycin. Details are as in panel A. The bottom graph shows results for sequences with a median enrichment in 5x puromycin that was at least two-fold higher than in 1x puromycin. D. Consensus sequences for 11mers that were at least five-fold more enriched with 1x puromycin, and either as enriched, or even more enriched, with 5x puromycin. Results for all 194 sequences are shown at the top, those for 37 sequences with at least one G and no 6mer PPT are at the bottom, and those for the majority that remained are in the middle.

After selection in 1 µg/ml puromycin, there were 230 different 11mers with median enrichment of at least five-fold. Further selection with 5 µg/ml gave 266 sequences. In both cases, there was selection for C and U residues and strong loss of G (Figure 4C). 194 sequences were five-fold enriched under both conditions, and at least as abundant with 5 µg/ml as with 1 µg/ml. These showed a preference for U or C in all 11 positions, with particularly strong preference for U at position -4 relative to the AG dinucleotide (Figure 4D). 80 of the 11mers included at least ten pyrimidines and 126 included at least nine, and this preference was strongest for sequences that increased in abundance under higher selective pressure (Figure 4C).

Strangely, 45 of the selected sequences lacked six consecutive pyrimidines, and included one or more G residue. These had a strong consensus sequence (Figure 4D, panel labelled “G no PPT”) which is completely absent from the reference genome. We also looked for some of the related sequences in the 50 nt upstream of all splice sites and found only very few examples, all of them upstream of a consensus U/C-rich PPT. To find out whether “CCCAGUGUGGC” was active, we also tested it individually, but cells containing the plasmid were as susceptible to puromycin as the wild-type (not shown). We therefore cannot explain why this consensus was selected in our screen.

### Tethering of CPSF3 or SF1 enhances splicing

Next, we tested the effect of expressing lambdaN-tagged proteins on mRNA processing (Figure 5A). We first used the reporter with (U)_3_ at the PPT position, aiming to identify factors that stimulated processing. In each case, the expressed protein had a lambdaN peptide at the N terminus, and two myc tags at the C-terminus. All results from the tethering must be interpreted with caution because protein function might be impaired or altered by the presence of the tags. Also, the tethered protein might have an inappropriate orientation or position relative to the RNA backbone or the splice site. As a negative control, we expressed lambdaN-GFP-2myc, which had no effect on processing or puromycin resistance (Figure 5B, replicates and Western blots in Supplementary Figure S1). Next, we tethered CPSF73, the cleavage and polyadenylation factor, and saw strongly enhanced processing at the “correct” sites and a more than ten-fold increase in puromycin resistance (Figure 5C, Supplementary Figure S2). This was surprising since we had expected this enzyme to cleave near the tethering site. Tethering of the splicing factor SF1 also enhanced processing (Figure 5D, Supplementary Figure S2). In contrast, tethering of the putative homologue of U2AF65 shunted some processing to the upstream positions, with marginally reduced puromycin resistance (Figure 5E, Supplementary Figure S3).

**Figure 5.**
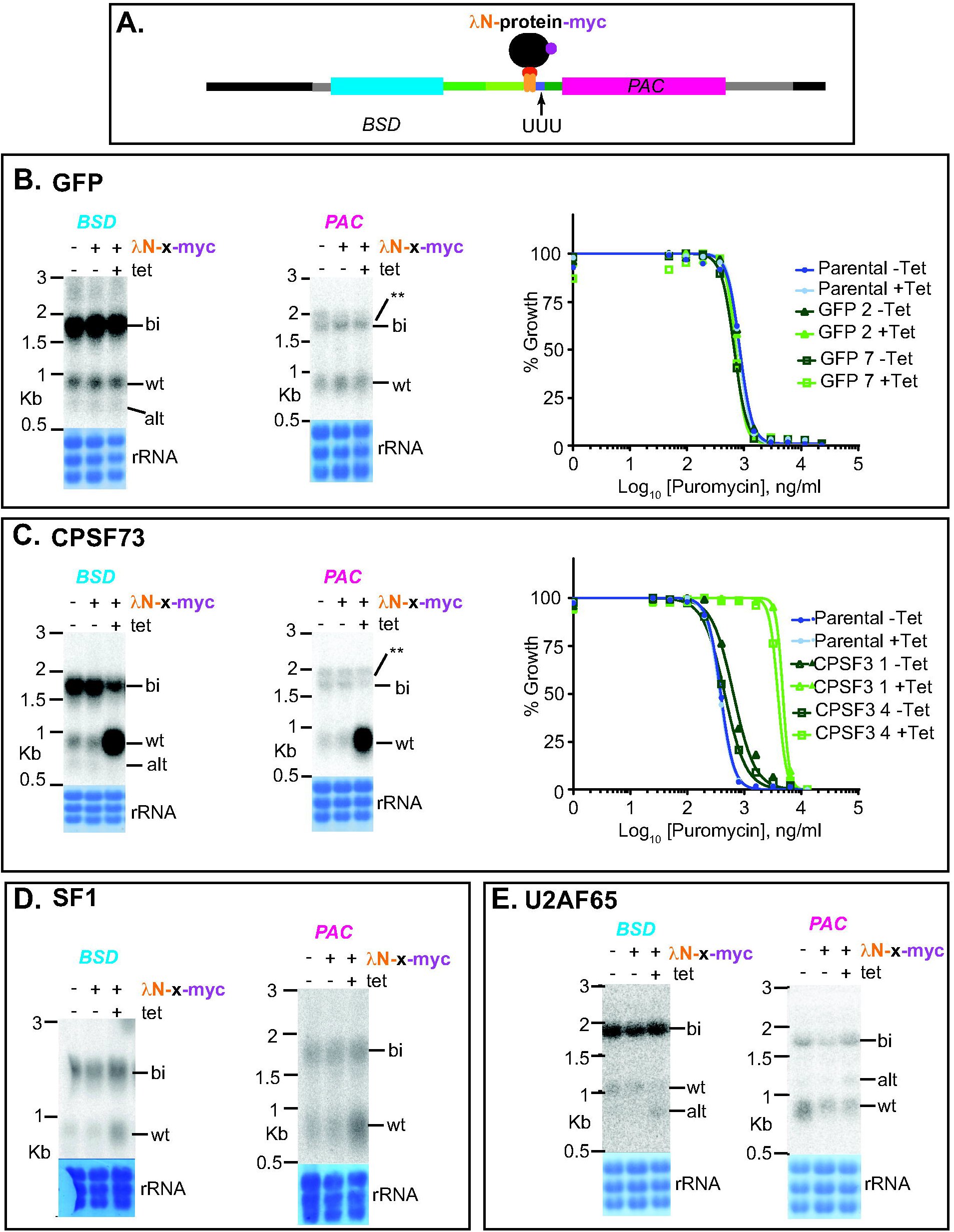
Effects of tethering GFP, CPSF3, SF1 or U2AF65 to the boxB-(U)_3_ reporter. A. Cartoon showing the precursor before processing, with tethered protein. B. Tethering of lambdaN-GFP-myc. In this and all subsequent similar panels, a Northern blot is shown for cells with no inducible lambdaN protein, then one clone with and without tetracycline. Results for both *BSD* and *PAC* are illustrated, with the stained rRNA on the membrane below. The susceptibility of two clones to puromycin is shown on the right. Other labels are as in Figure 1C. Results for additional clones and biological replicates are shown in Supplementary Figures S1-S3. C. Tethering of lambdaN-CPSF73. D. Tethering of lambdaN-SF1. E. Tethering of lambdaN-U2AF65.

### Tethering of potential splicing regulators

Next, we investigated nuclear proteins that had previously been shown to influence mRNA levels. DRBD3 and DRBD4 are possible homologues of Opisthokont PTBs. Tethering of either protein inhibited processing between *BSD* and *PAC* entirely, with increased puromycin susceptibility (Figure 6A, B, Supplementary Figure S4). In contrast, tethering of HNRNPF/H had no effect (Figure 6C, Supplementary Figure S1). Tethering of the SR-domain protein TSR1 and its interaction partner TSR1IP, in contrast, resulted in use of the alternative sites (Figure 7A, B; Supplementary Figure S5), with decreased puromycin resistance. We also investigated two additional SR-domain proteins: RBSR1, which is mainly in the nucleus, and RBSR2, which is present in both nucleus and cytoplasm (Dean et al. 2016). Each of these had the same effect as TSR1 (Figure 7C,D; Supplementary Figure S6).

**Figure 6.**
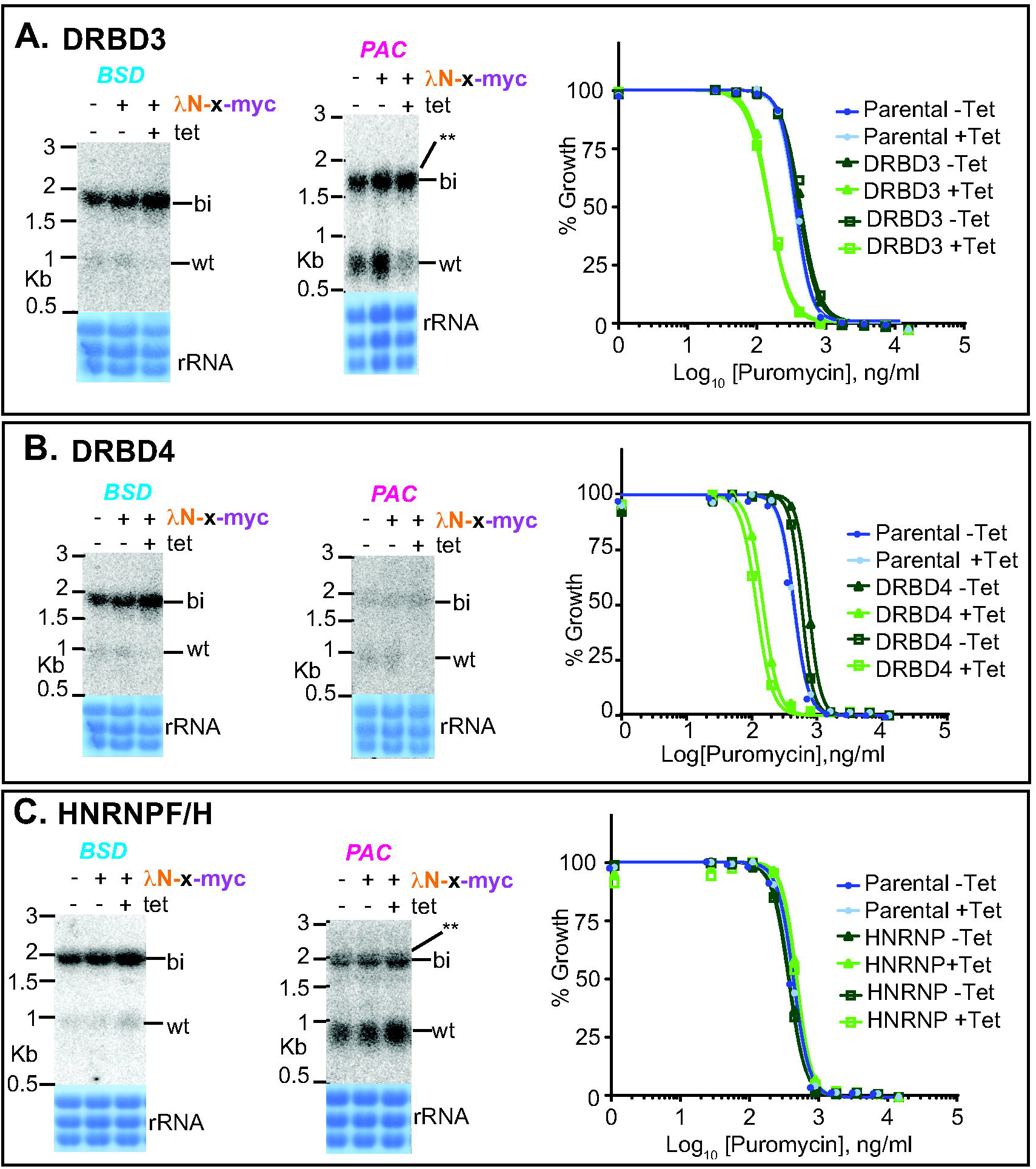
Effects of tethering (A) DRBD3, (B) DRBD4 or (C) HNRNPF/H to the boxB-(U)_3_ reporter. Details are as for Figure 5. See also Supplementary Figures S1 and S4.

**Figure 7.**
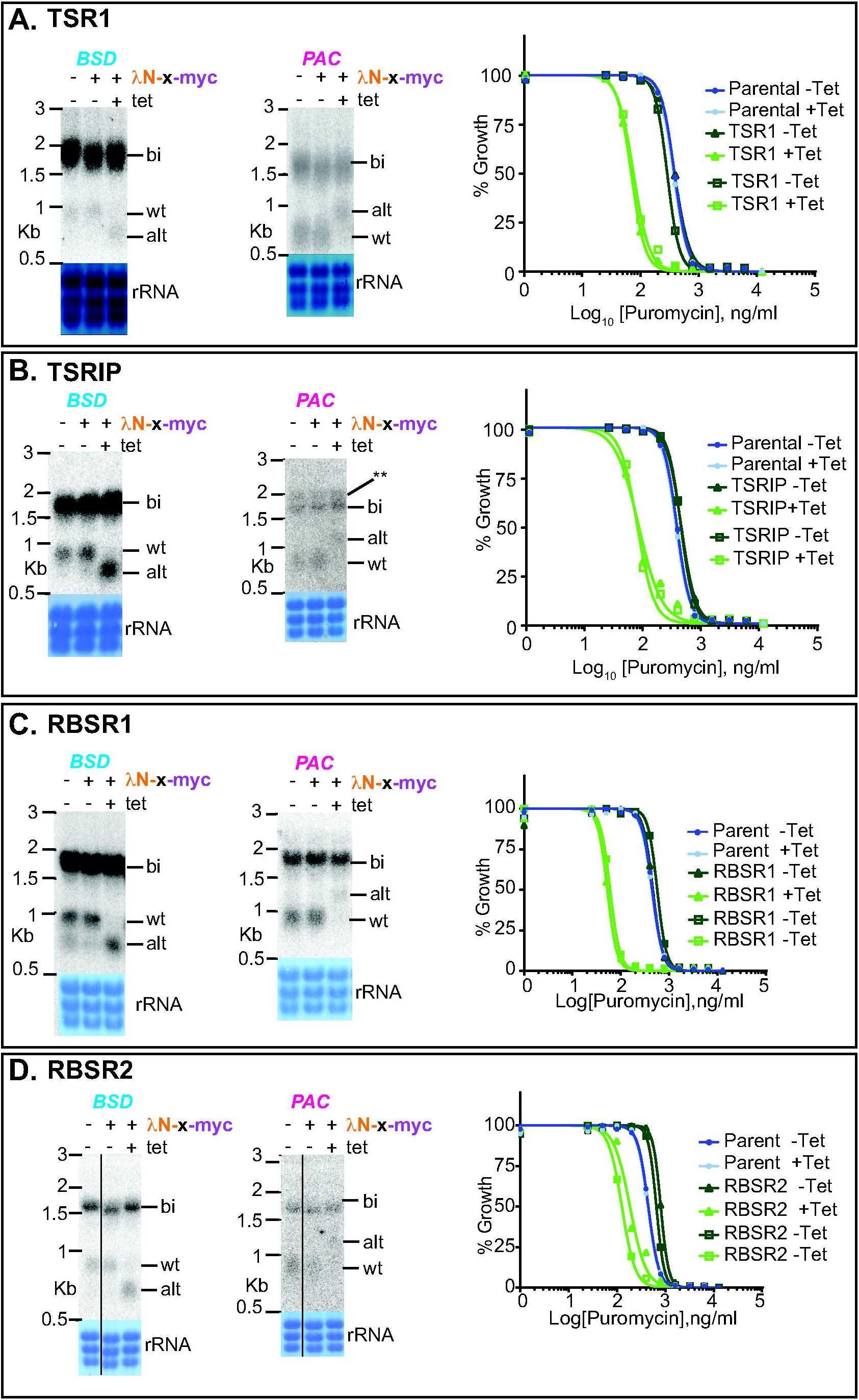
Effects of tethering SR-domain proteins to the boxB-(U)_3_ reporter. Details are as for Figure 5. See also Supplementary Figures S5 and S6.

So far, the experiments had employed a reporter with an extremely short PPT. We therefore asked whether the potential regulators would also affect splicing of a precursor with stronger PPTs. Using the 2xboxB-(U)_9_ reporter, HNRNPF/H and U2AF65 again had no effect. DRBD3 and DRBD4 inhibited splicing, and the SR-domain proteins caused use of the alternative sites (Figure 8; Supplementary Figures S7-S9). With 2xboxB-(U)_14_, effects were similar except that for RBSR1 and RBSR2, splicing was inhibited and the alternatively processed products were not detected (Figure 8; Supplementary Figures S7-S9).

**Figure 8.**
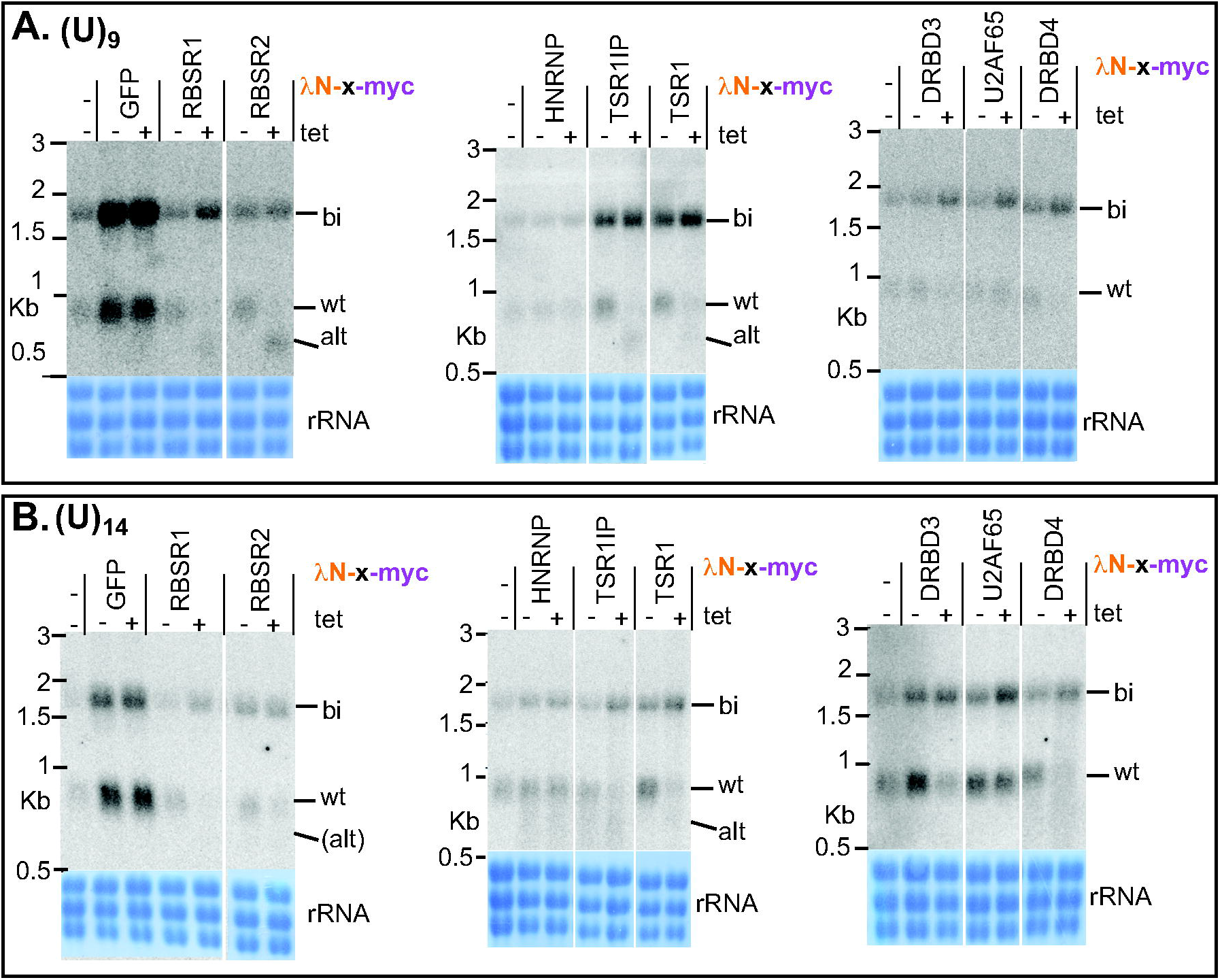
Effects of tethering different proteins to boxB-(U)_9_ and boxB-(U)_14_ reporters. Northern blots are shown for the *BSD* mRNAs. Additional clones and replicates are shown in Supplementary Figures S7-S9.

### The protein associations of RBSR1 and RBSR2

Since RBSR1 and RBSR2 had not previously been investigated in *T. brucei*, we characterized their protein associations. Cell lines were made that expressed C-terminally tandem affinity-tagged RBSR1 or RBSR2 from the endogenous locus. After purification over IgG beads, the tagged proteins were released using His-tagged TEV protease, which was subsequently removed using a nickel column. The extracts were then analysed by mass spectrometry, using inducibly-expressed, similarly tagged GFP as the control. RBSR1 associated with 131 proteins, including 22 components of the splicing machinery (Figure 9A, Supplementary Table S3), five components of the PRP19 complex as well as U2 and U5 snRNP components (Supplementary Table S4). Interestingly, the CFII subunit of the polyadenylation complex was also present. Although RBSR1 is predominantly in the nucleus, it was also associated with ribosomes and with one of the cap-binding translation initiation complexes, EIF4E4/EIF4G3. There was little overlap with proteins that co-immunoprecipitated with GFP-tagged *Trypanosoma cruzi* RBSR1, although that study also identified TRRM1 and some ribosomal proteins (Wippel et al. 2018). Our study was probably more sensitive since we used quantitative proteomics.

**Figure 9.**
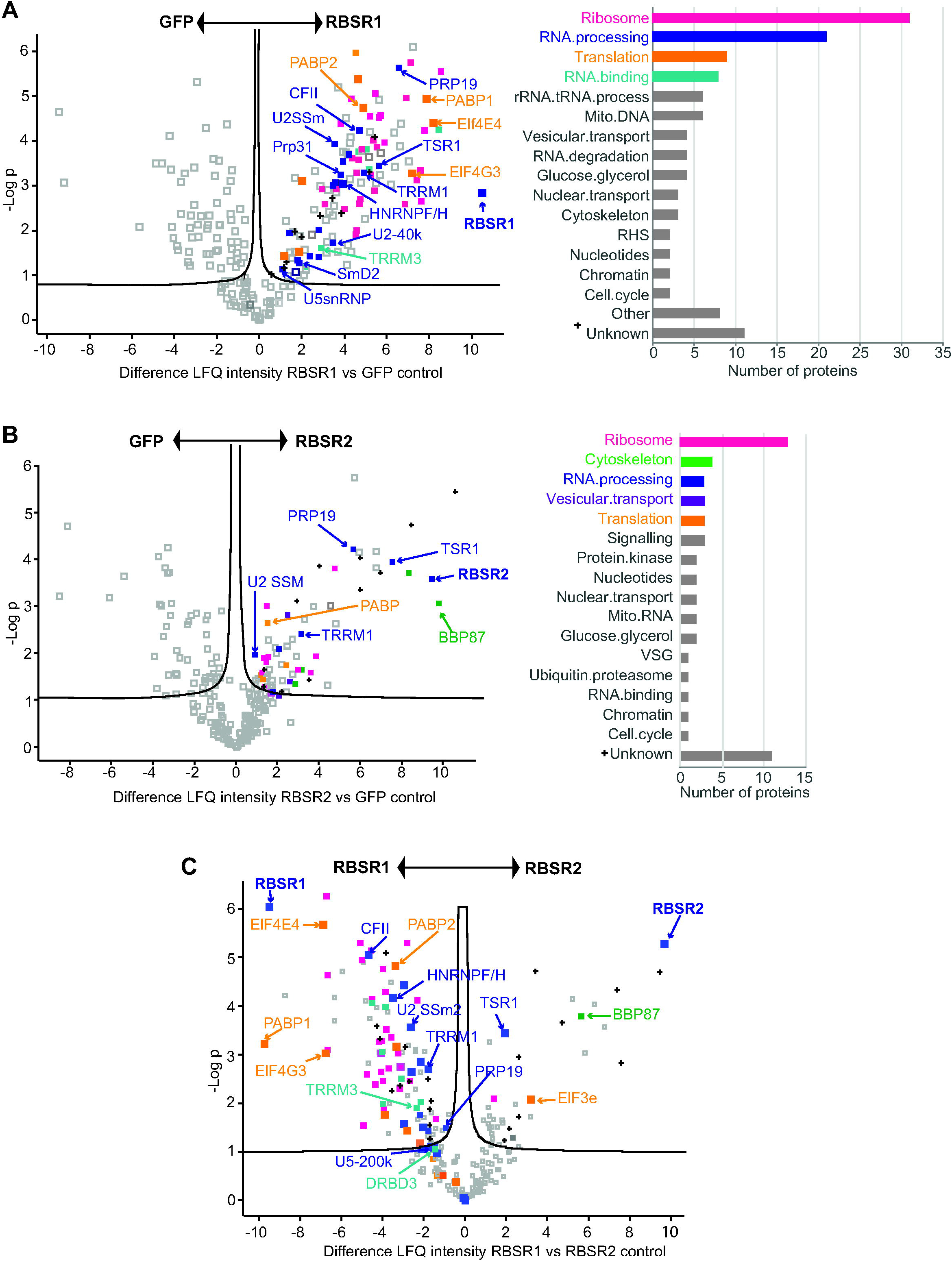
Proteins associated with RBSR1 and RBSR2. RBSR1-TAP and RBSR2-TAP were purified on IgG columns and released with TEV protease. Triplicate preparations were analyzed by mass spectrometry, using GFP-TAP as the control. Results were generated by PERSEUS. Proteins of interest are indicated. Functional categories were assigned manually from the annotation (see Supplementary Table S3). A. RBSR1 compared with GFP. Numbers of enriched proteins in particular categories are shown on the right. B. RBSR2 compared with GFP. Numbers of enriched proteins in particular categories are shown on the right. C. RBSR2 compared with RBSR1.

The RBSR2 purification yielded fewer (56) specifically associated proteins (Figure 9B, Supplementary Table S3), perhaps because RBSR2 is (according to quantitative mass spectrometry) about four-fold less abundant (Tinti and Ferguson 2022) (Supplementary Table S4). RBSR2 was not clearly associated with the mRNA processing machinery but pulled down four proteins from the basal body, and again various ribosomal proteins (Figure 9B, Supplementary Table S3). A comparison of the two proteomes (Figure 9C) revealed that although TSR1 was present in both, it was more enriched with RBSR2. Vault proteins copurified with both, but more with RBSR2, and RBSR1 pulled down many more (potentially) regulatory RNA-binding proteins, including DRBD3.

### Effects of RBSR1 and RBSR2 depletion on growth and the transcriptome

To examine the roles of RBSR1 and RBSR2 in mRNA metabolism, we depleted them by inducible RNA interference (RNAi) and examined the transcriptomes, starting with cells expressing the corresponding TAP-tagged protein. The lines with best depletion of RBSR1 had, without induction, roughly half as much RBSR1 protein as the starting (tagged) cell line. After RNAi induction, the level decreased to about 20% (Figure 10A, Supplementary Figure S10). The cell lines grew almost normally and RNAi induction had no effect (Figure 10A). For RBSR2, RNAi decreased the protein level to around 11%, but there was also no effect on growth (Figure 10B). We compared the transcriptomes of two independent *RBSR1* RNAi lines and three independent *RBSR2* RNAi lines (Figure 10A, Supplementary Figure S10) with three replicates from the RBSR2-TAP starting line (chosen because the TAP tag alone had no effect on growth). For both RBSR1 and RBSR2, RNAi induction had no effect on the transcriptome (Supplementary Figure S8, Supplementary Tables S5 and S6). All of the RBSR RNAi lines showed accumulation of the RNAi-mediating RNA. In the sense direction, examination of the reads (not shown) revealed *trans*-splicing of the “sense” strand downstream of the normal initiation codon. We pooled the results with and without tetracycline, since it made no difference, and compared the transcriptomes of the two lines with the starting (TAP-tagged) line for RBSR2 (labelled as “WT” in the Figures and Tables). Both RNAi lines had increased expression of variant surface glycoprotein genes that are normally silent, which might be a sign of stress. For both poly(A)+ (Figure 10C) and rRNA-depleted RNA (Figure 10D), the RBSR2 line showed preferential accumulation of shorter RNAs and loss of longer ones, but hardly any differences exceeded two-fold. The amount of mRNA, as judged by hybridization of a Northern blot with a spliced leader probe, was also twice that in the starting cell line. Since this effect is unrelated to the RBSR2 level and also to the cell density (Supplementary Figure S8), we cannot relate it to RBSR2 function. For RBSR1, we also noted small, mostly less than two-fold differences compared to the same “WT” cell line. For poly(A)+ mRNA, there was preferential loss of longer RNA (Figure 10E) but the correlation was weaker than for RBSR2, and absent for rRNA-depleted RNA (Figure 10F). Detailed examination of the read alignments for several genes that showed such discrepancies revealed no changes in polyadenylation or splice sites.

**Figure 10.**
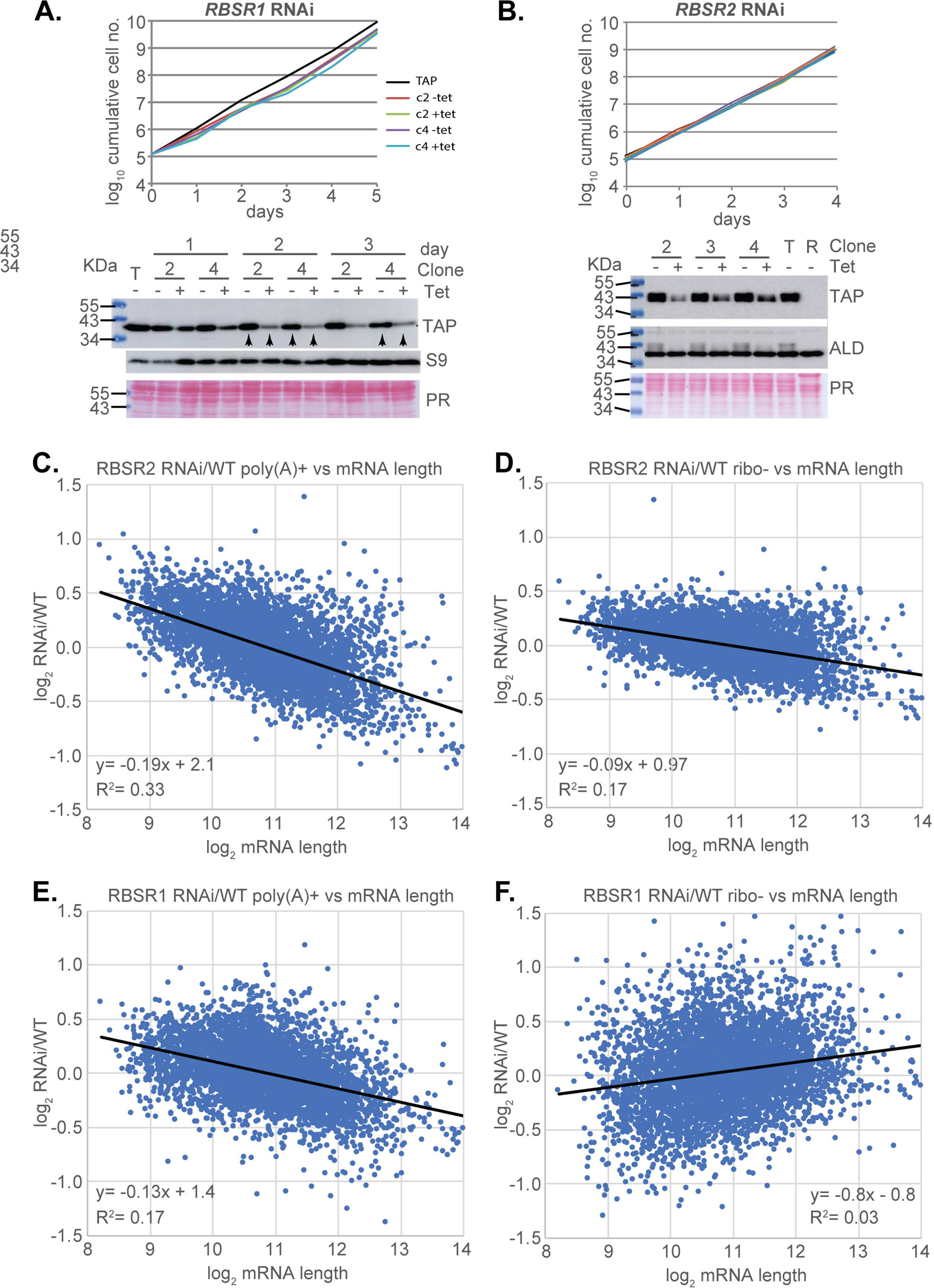
Transcriptomes of trypanosomes containing integrated plasmids for RBSR1 and RBSR2 RNAi. A. Growth of two clones (c2 and c4) with and without *RBSR1* RNAi induction. Cells with the tagged protein (TAP) are shown for comparison, and the Western blot is shown below. Further (uncropped) results are shown in Supplementary Figure S11. The arrows indicate the samples taken for RNASeq. B. Growth of two clones with and without *RBSR2* RNAi induction. Cells with the tagged protein (TAP) are shown for comparison, and the Western blot is shown below. The lines are not labelled because they are indistinguishable. Further (uncropped) results are shown in Supplementary Figure S12. C. Poly(A)+ transcriptomes from the RBSR2 RNAi line, with and without tetracycline, compared with the “wild-type” (cells with tagged RBSR2). The ratio of RBSR2 line/WT is shown on the y-axis and the mRNA length on the x-axis. The formula for the regression line and the Pearson correlation coefficient are also shown. For more details, see Supplementary Figures S12 and S13, as well as Supplementary Table S6. D. rRNA-depleted transcriptomes from the RBSR2 RNAi line, with and without tetracycline, compared with the “wild-type” (cells with tagged RBSR2). The ratio of RBSR2 line/WT is shown on the y-axis and the mRNA length on the x-axis. E. Poly(A)+ transcriptomes from the RBSR1 RNAi line, with and without tetracycline, compared with the “wild-type” (again, cells with tagged RBSR2, since the tagging had no effect on growth). The ratio of RBSR1 line/WT is shown on the y-axis and the mRNA length on the x-axis. D. rRNA-depleted transcriptomes from the RBSR1 RNAi line, with and without tetracycline, compared with the “wild-type” (cells with tagged RBSR2). The ratio of RBSR1 line/WT is shown on the y-axis and the mRNA length on the x-axis.

## Discussion

The results in this paper confirm that a wide range of different PPTs can be used to specify *trans*-splicing acceptor sites in trypanosomes. There was a slight preference for U over C, which was somewhat less pronounced than is seen in comparisons of sites used *in vivo* (Kolev et al. 2010). The effects of including two boxB stem-loops suggested that the presence of a secondary structure could influence splice site choice. A study in yeast revealed that intron secondary structures can indeed influence splicing kinetics (Barrass et al. 2015). It is also possible that the two boxBs affect DNA structure and slow down RNA polymerase II elongation, enhancing splicing. In trypanosomes, RNA polymerase II tends to accumulate in regions preceding splice acceptor sites, perhaps indicating an elongation delay (Wedel et al. 2017).

The results of our tethering experiments suggest that this will be a useful approach both to investigate the functions of putative splicing factors, as well as to screen for both repressors and activators. As expected, tethering of SF1 stimulated splicing at the “wild-type” site. We had predicted that tethering of CPSF73 would cause aberrant polyadenylation, but instead, it also stimulated “correct” splicing. Presumably tethered CPSF73 acted by recruiting splicing factors as well as the rest of the polyadenylation complex, but it is not clear why the “wild-type” acceptor was then used. Surprisingly, tethering of U2AF65 to the (U)_3_ reporter weakly promoted alternative splicing, perhaps also stabilizing the bicistronic RNA. With the (U)_9_ and (U)_14_ reporters, no more alternative splicing was seen but the bicistron was again more abundant. Depletion of trypanosome U2AF65 inhibits splicing, and it interacts with SF1 but not U2AF35 (Vazquez et al. 2009). There is, however, evidence that it can stabilize a bound mRNA. Perhaps in this case, the tethered protein was impaired in splicing function due to the presence of the tags, or because of inappropriate orientation relative to the RNA substrate.

DRBD3 and DRBD4 (De Gaudenzi et al. 2006; Estévez 2008; Stern et al. 2009) are the closest homologues of mammalian PTB (polypyrimidine-tract-binding) proteins, which have four RNA-recognition motifs (RRMs) (Keppetipola et al. 2012). DRBD3 has two RRMs, and a glutamine-rich disordered region at the C-terminus. DRBD4 has four RRM domains, although one is rather weak (Stern et al. 2009). DRBD4 is predominantly found in the nucleus, whereas DRBD3 is slightly concentrated in the nucleus but also in the cytoplasm. RNAi targeting either DRBD3 or DRBD4 was shown to reduce SLRNA levels but this could have been a secondary effect of growth inhibition (Stern et al. 2009). RNAi against either DRBD3 or DRBD4 causes both increases and decreases in transcript levels, some of which can be attributed to changes in mRNA stability (Estévez 2008; Stern et al. 2009). A conserved binding motif, UUCCCCUCU, was identified for DRBD3 (Das et al. 2015) and found in the 3’-UTRs of nearly 300 different mRNAs (Gupta et al. 2013; Das et al. 2015). In our screen, this sequence was present in a single 11mer, but the read count numbers were too low to assess enrichment after selection.

Cumulative evidence suggests that mammalian PTB represses intron inclusion by binding either within the exon, or at the 3’ splice site. For weak splice sites, a single binding site may be sufficient, whereas for efficient ones, two sites are needed (Keppetipola et al. 2012). Suggested mechanisms of action including looping out the branch-point adenosine, and looping out entire exons through interaction with upstream and downstream PPTs (Romanelli et al. 2013). Both DRBD3 and DRBD4 indeed repressed *PAC trans* splicing when tethered, even when the reporter included a strong PPT. The consensus binding motif for DRBD3 is not present in the intergenic region, so attachment via the high-affinity lambdaN-boxB interaction may be sufficient. Still, interactions with the weak splicing signal in the intergenic region, or with downstream PPTs, are also possible. The sequence CCUCCCCUUCU is at the end of the *PAC* open reading frame. DRBD3 pulled down U1-70K and U1C, consistent with a role in splicing (Fernandez-Moya et al. 2012), but this would suggest recruitment of splicing factors, rather than repression. Some effects of either DRBD3 or DRBD4 might possibly be due to stabilization of the bicistronic precursor. DRBD3 is associated with the two poly(A)-binding proteins PABP1 and PABP2 (Fernandez-Moya et al. 2012), consistent with roles in stabilizing cytoplasmic mRNAs. In addition, DRBD3 is associated with a protein (Tb927.9.4080) with putative capping activity that is associated with the putative translation initiation complex containing EIF4E5 and EIF4G1 (Fernandez-Moya et al. 2012; Freire et al. 2014).

In Opisthokonts, SR-domain splicing regulators are known to define exon junctions during *cis*-splicing by binding at the beginning of the exon, which in most cases is within the coding region (Howard and Sanford 2015). Although some Kinetoplastid SR-domain proteins had previously been investigated (Gupta et al. 2014; Levy et al. 2015; Wippel et al. 2018), their functions were unknown. We found that tethering TSR1 and its interaction partner TSR1IP shifted splicing to the alternative upstream location even when a strong PPT was present downstream of the tethered protein. This is consistent with these proteins also acting to define the start of an exon. RBSR1 and RBSR2 shifted splicing upstream when the (U)_3_ or (U)_9_ reporters were used. With the (U)_14_ reporter, use of the “wild-type” acceptor site was inhibited but the alternative products were not detected.

Since RBSR1 and RBSR2 had not previously been specifically investigated, we characterized their protein interactomes and the effects of depletion. Depletion of either protein had little effect on cell proliferation, and transcriptome differences relative to a line without RNAi were unrelated to the level of the target protein. A clear discrepancy between poly(A)+ and rRNA-depleted RNA in the cell line with RBSR1 RNAi was intriguing but difficult to interpret. If it was caused by the 50% loss of RBSR1, why was it not more severe after RNAi induction? One possibility was that selection of the RNAi lines (two or three independent clones) had by itself caused transcriptome changes. These issues require further investigation. Meanwhile, the proteins associated with RBSR1 and RBSR2 were much more instructive. RBSR1 preferentially copurified proteins associated with splicing, most particularly five components of the PRP19 complex, which is thought to associate with the spliceosome when the U4 snRNP dissociates from U5 and U6. This, and the presence of two U2 components, is consistent with RBSR1 affecting splice acceptor site choice. Intriguingly, RBSR1 also specifically pulled down only one of the cap-binding translation initiation factor complexes, EIF4E4 and EIF4G3, even though two others (EIF4E3/EIF4G4 and EIF4E6/EIF4G5) are as abundant (Tinti and Ferguson 2022), and RBSR1 is detected exclusively in the nucleus. Results from *Leishmania* suggest that the EIF4E4/G3 complex is implicated in translation of mRNAs encoding ribosomal proteins (Assis et al. 2021). This link, as well as the apparent association of RBSR1 with the ribosome, might merit further investigation.

RBSR2 was associated with only three splicing-related proteins: TSR1, TRRM1 and PRP19. It is thus possible that the effect of tethered RBSR1 on splice site choice is due to its association with TSR1. Intriguingly, however, we also detected specific associations with proteins of the basal body. Inspection of available images on the tryptag.org web site (Dean et al. 2016) shows that in addition to predominantly nuclear localization, GFP-tagged RBSR2 gives diffuse cytoplasmic fluorescence and sometimes also one or two spots, but their location is probably not consistent with basal body association. RBSR2 also pulled down the three major vault proteins, which are related but not identical and all cytoplasmic when GFP-tagged (Dean et al. 2016). The function of vault proteins in trypanosomes is unknown.

Our results show that tethering of potential splicing regulators upstream of a *trans* splice acceptor site can give insights into factor function. The system designed here could also be used to screen for novel splicing factors with either stimulating or inhibitory functions. We expect that similar systems could be used in other organisms as well for analysis of both *cis* and *trans* splicing.

## Supplementary Material

Supplementary Text 1: Intergenic sequences.

Supplementary Table S1: Plasmids and oligonucleotides

Supplementary Table S2: 11mer sequences

Supplementary Table S3: Mass spectrometry of purified RBSR1-TAP and RBSR2-TAP.

Supplementary Table S4: Abundances of various processing-related proteins according to (Tinti and Ferguson 2022).

Supplementary Table S5: Mapped read counts for RBSR1 and RBSR2 RNAi lines and the cells expressing RBSR2-TAP.

Supplementary Table S6: DeSeq2 analysis of the results in Supplementary Table S5.

Supplementary Sequence 1: Ape file, pHD3180

Supplementary Sequence 2: Ape file, pHD3186

Supplementary Sequence 3: Ape file, pHD3190

Supplementary Sequence 4: Ape file, pHD3259

## Data

DNA sequencing data are available at Array express with accession numbers E-MTAB-11647 (preliminary PPT screen); E-MTAB-11628 (2nd PPT screen); E-MTAB-11627 (RBSR1 RNAi); and E-MTAB-11648 (RNSR2 RNAi).

## Acknowledgements

We thank Claudia Helbig and Ute Leibfried for technical assistance, and Claudia Helbig especially for several of the Northern blots. We also thank David Ibberson of the BioQuant sequencing facility (University of Heidelberg) for DNA library construction and sequencing. We thank Shula Michaeli for many useful discussions and Andreas Kulozik and Susanne Kramer for suggestions during advisory meetings. We are indebted to Nina Papavasiliou (DKFZ, University of Heidelberg) and Luise Krauth-Siegel (BZH, University of Heidelberg) for allowing us to share their laboratories including equipment and reagents after the flood in the ZMBH.

## Author contributions

AW performed the experiments, and LN directly supervised the initial work. AW and CC were both involved in conceptualization, methodology, data curation, formal analysis, validation, investigation and visualization, and writing the paper. OM analysed the sequencing data, supervised by DG. CC was responsible for overall supervision, funding acquisition, and project administration.

## Ethical standards

Not applicable.

