## Supplementary figures and images for "Sequences and proteins that influence mRNA processing in *Trypanosoma brucei*: evolutionary conservation of SR-domain and PTB protein functions"

### Supplementary Figure S1

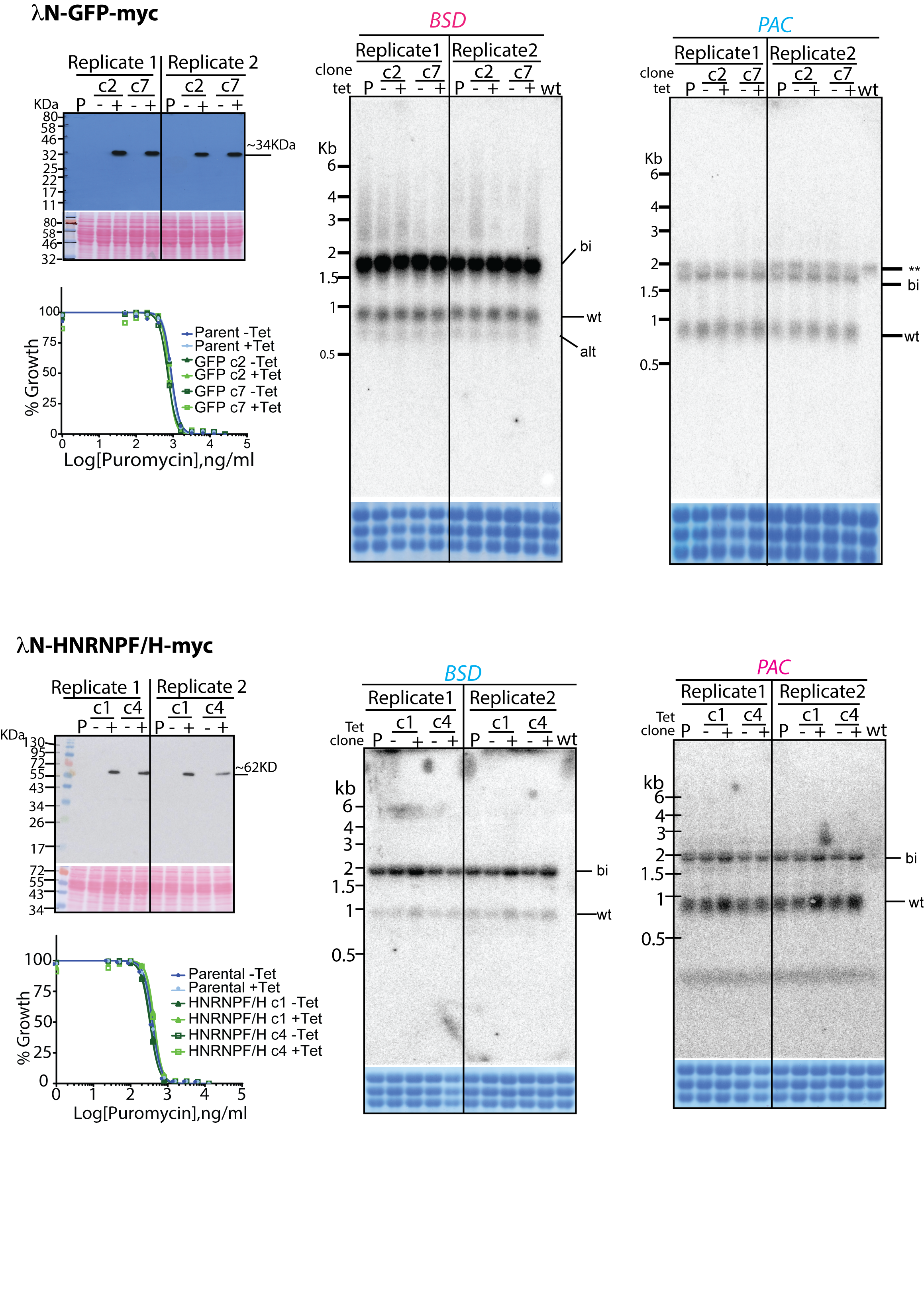

### Supplementary Figure S2

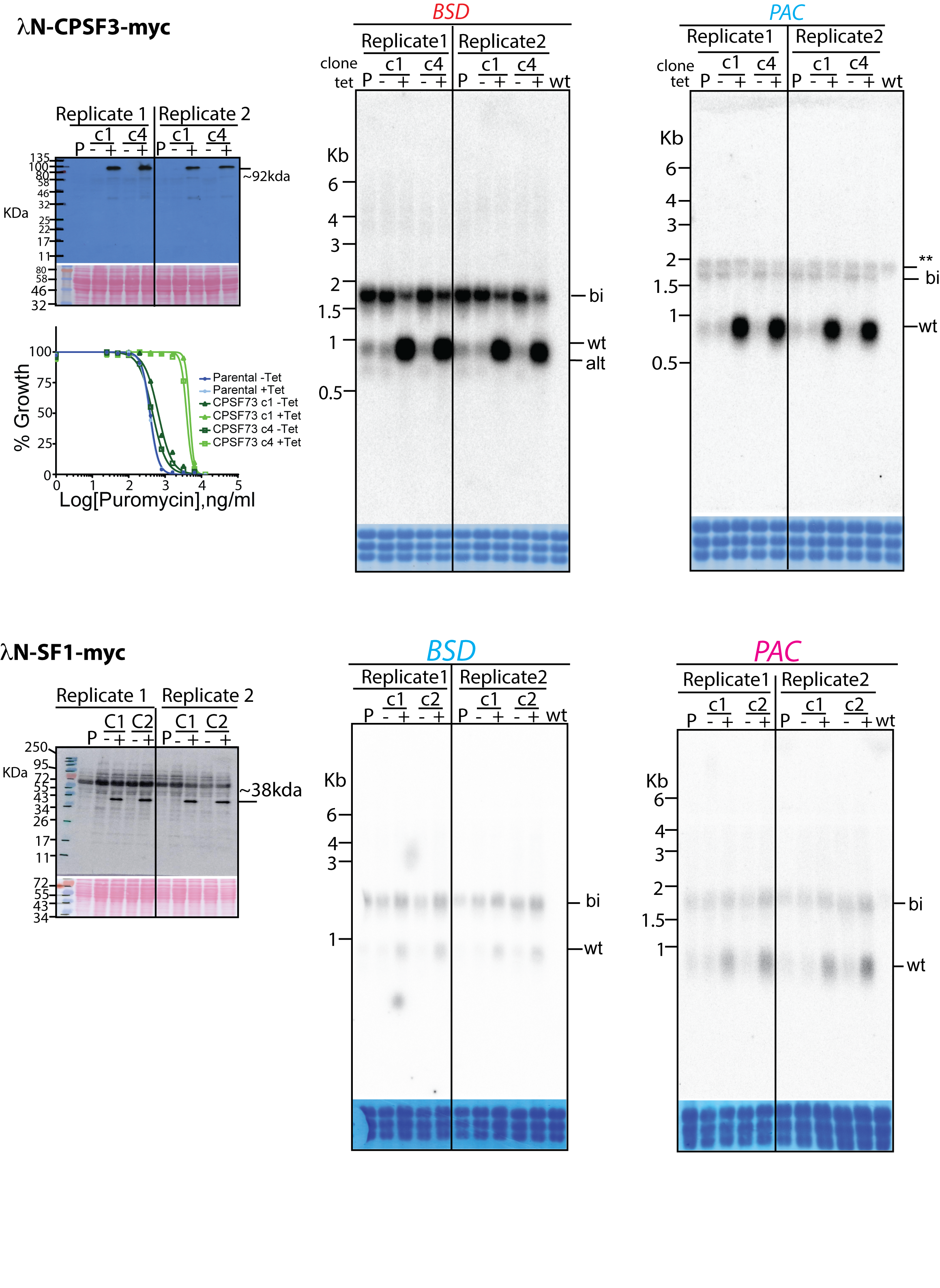

### Supplementary Figure S3

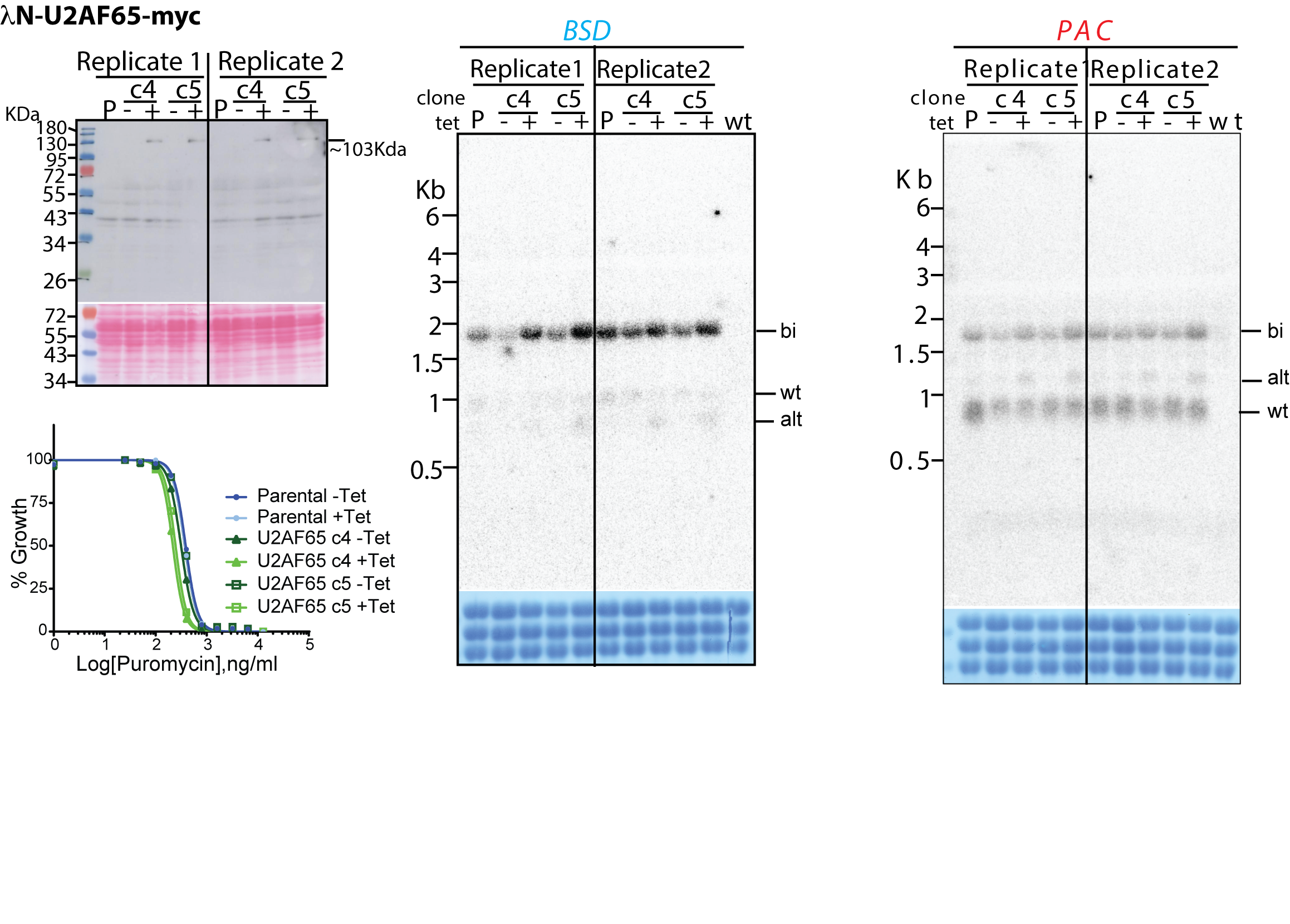

### Supplementary Figure S4

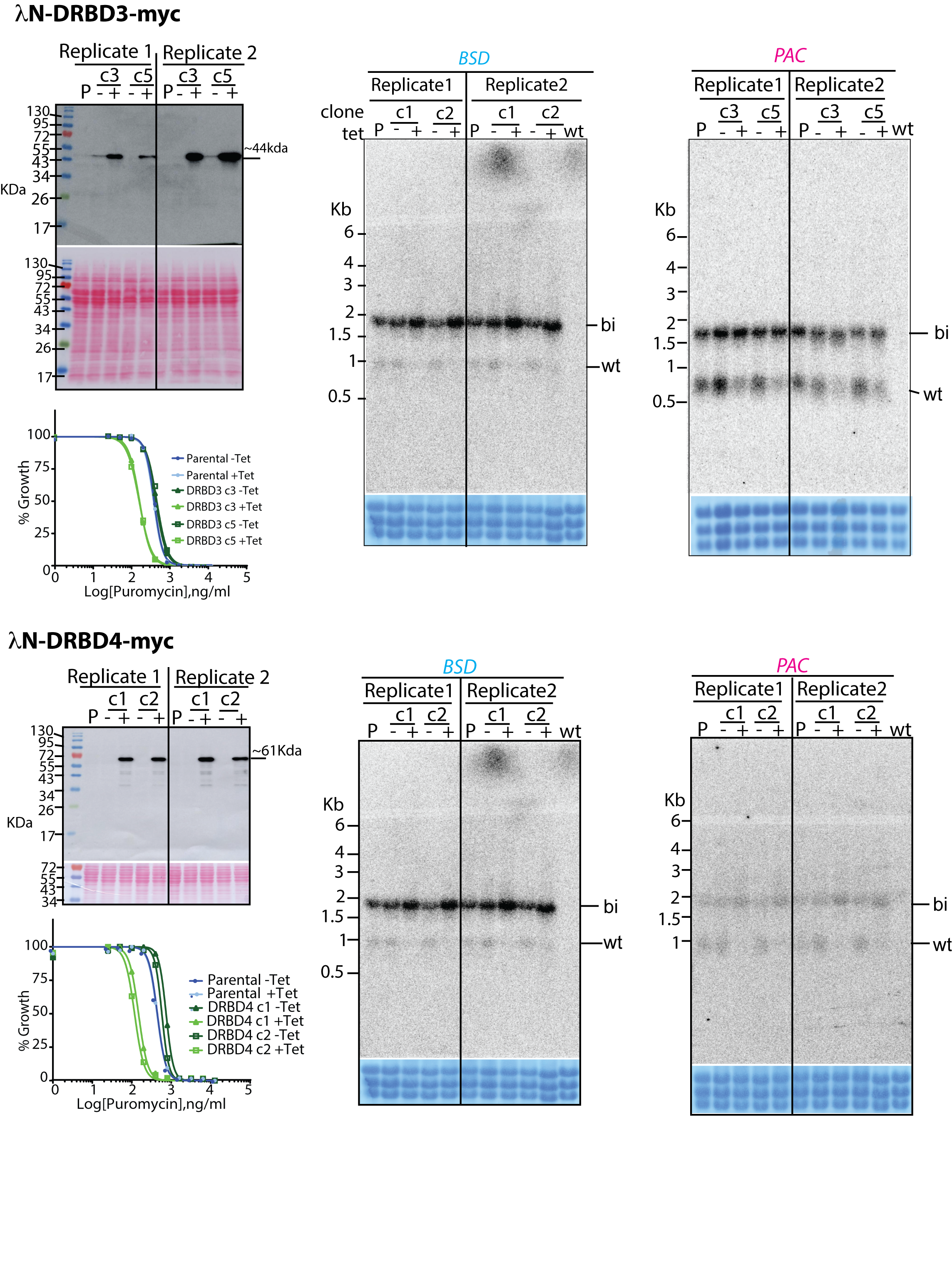

### Supplementary Figure S5

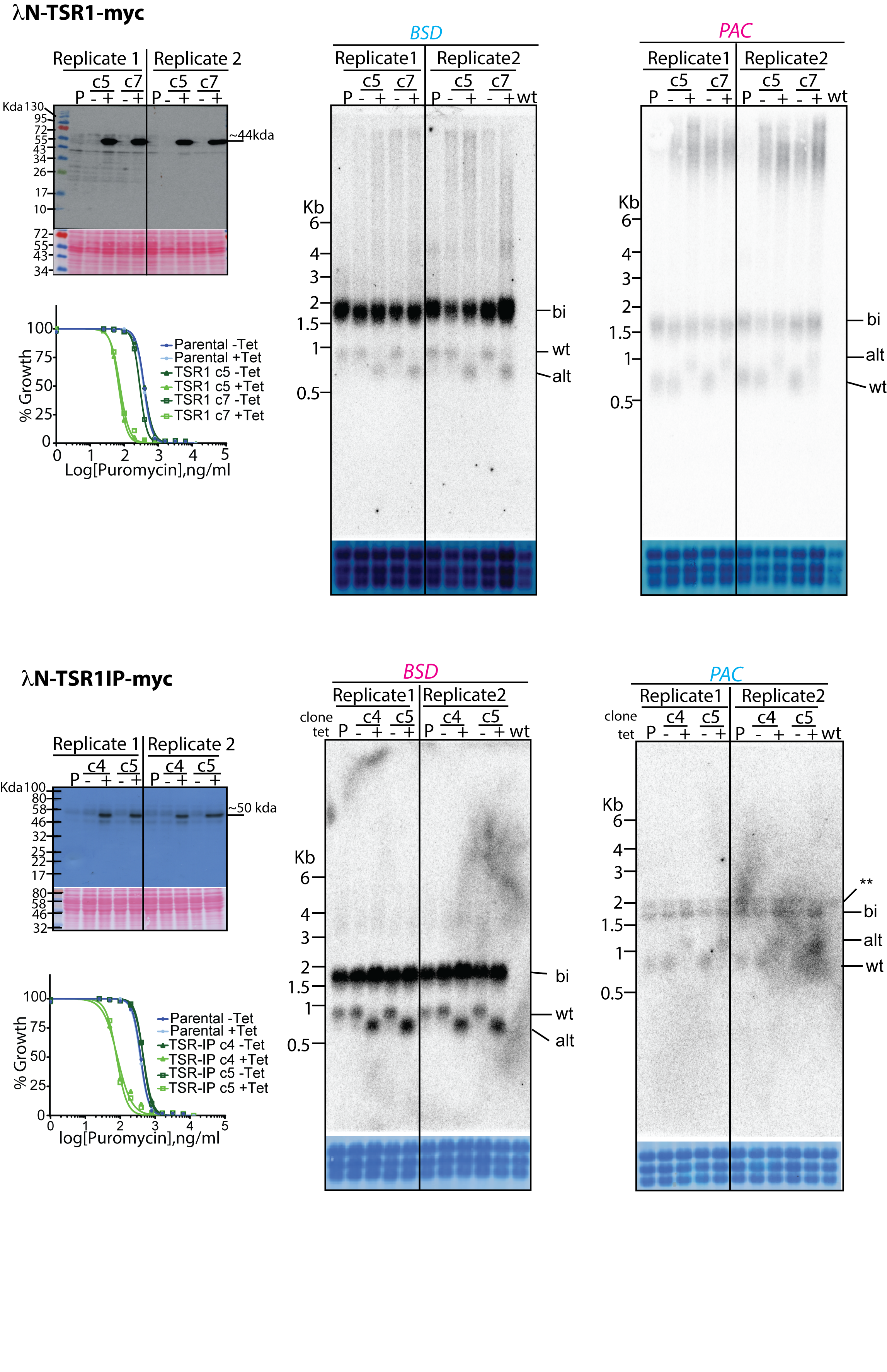

### Supplementary Figure S6

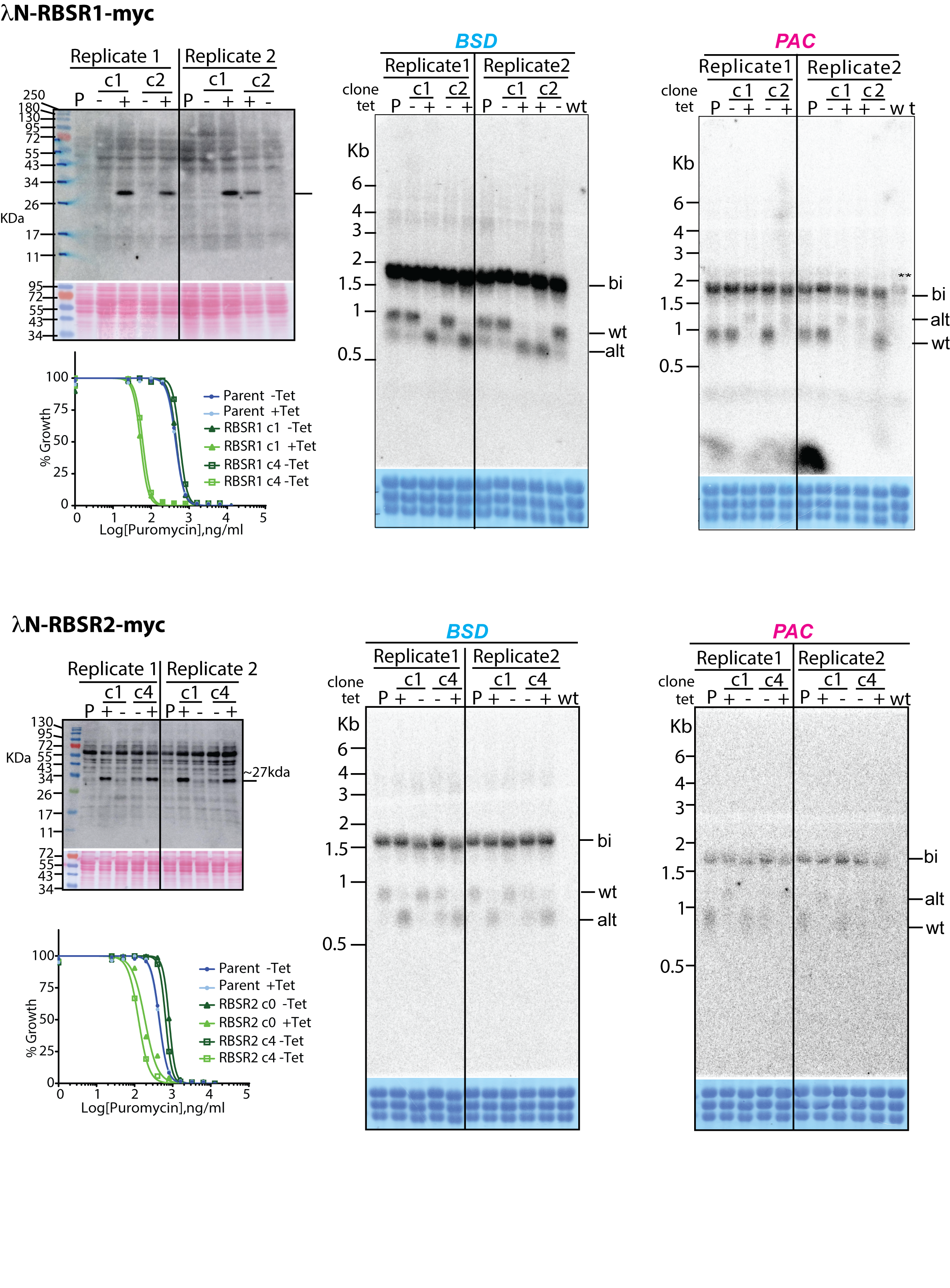

### Supplementary Figure S7

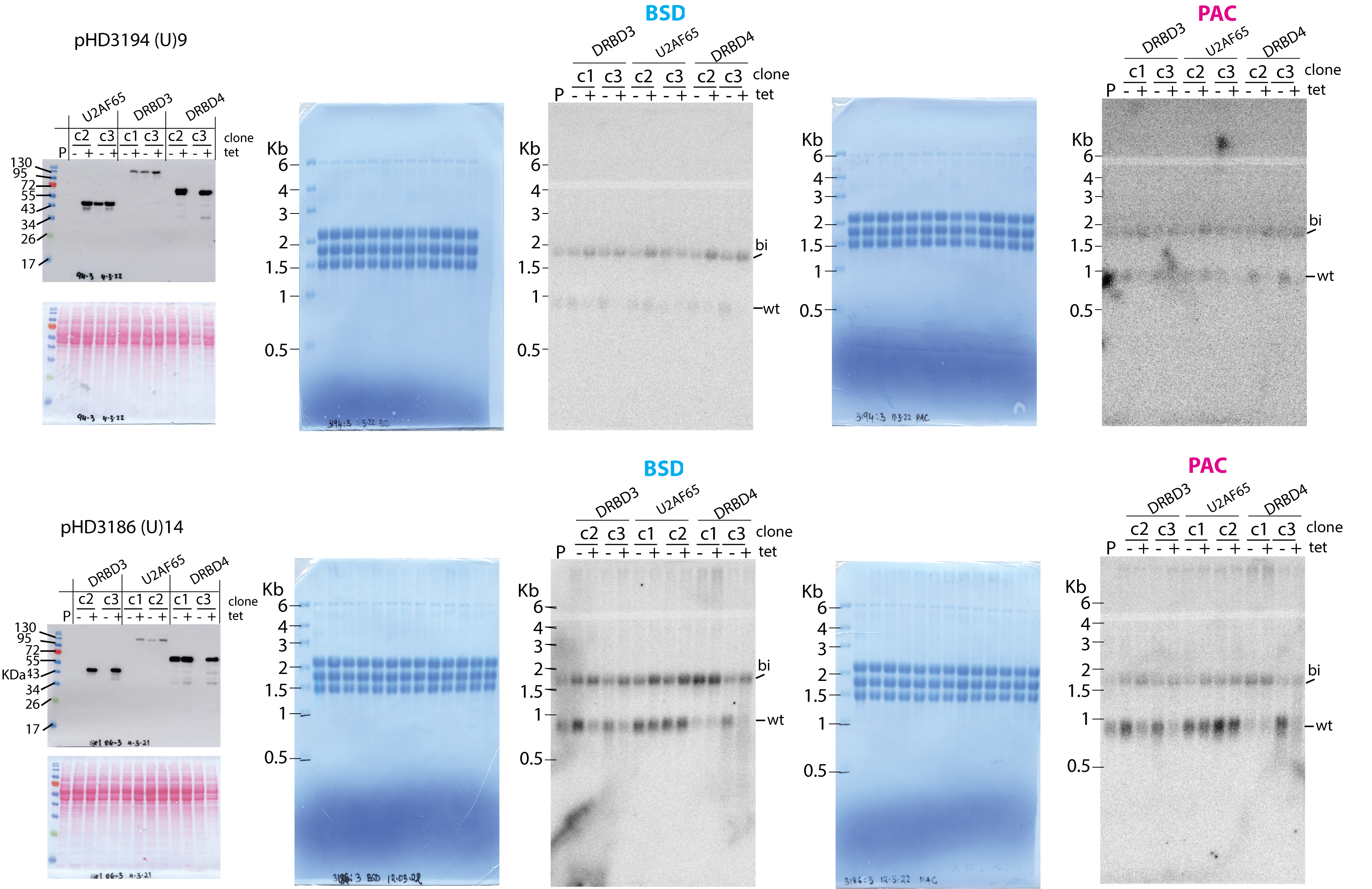

### Supplementary Figure S8

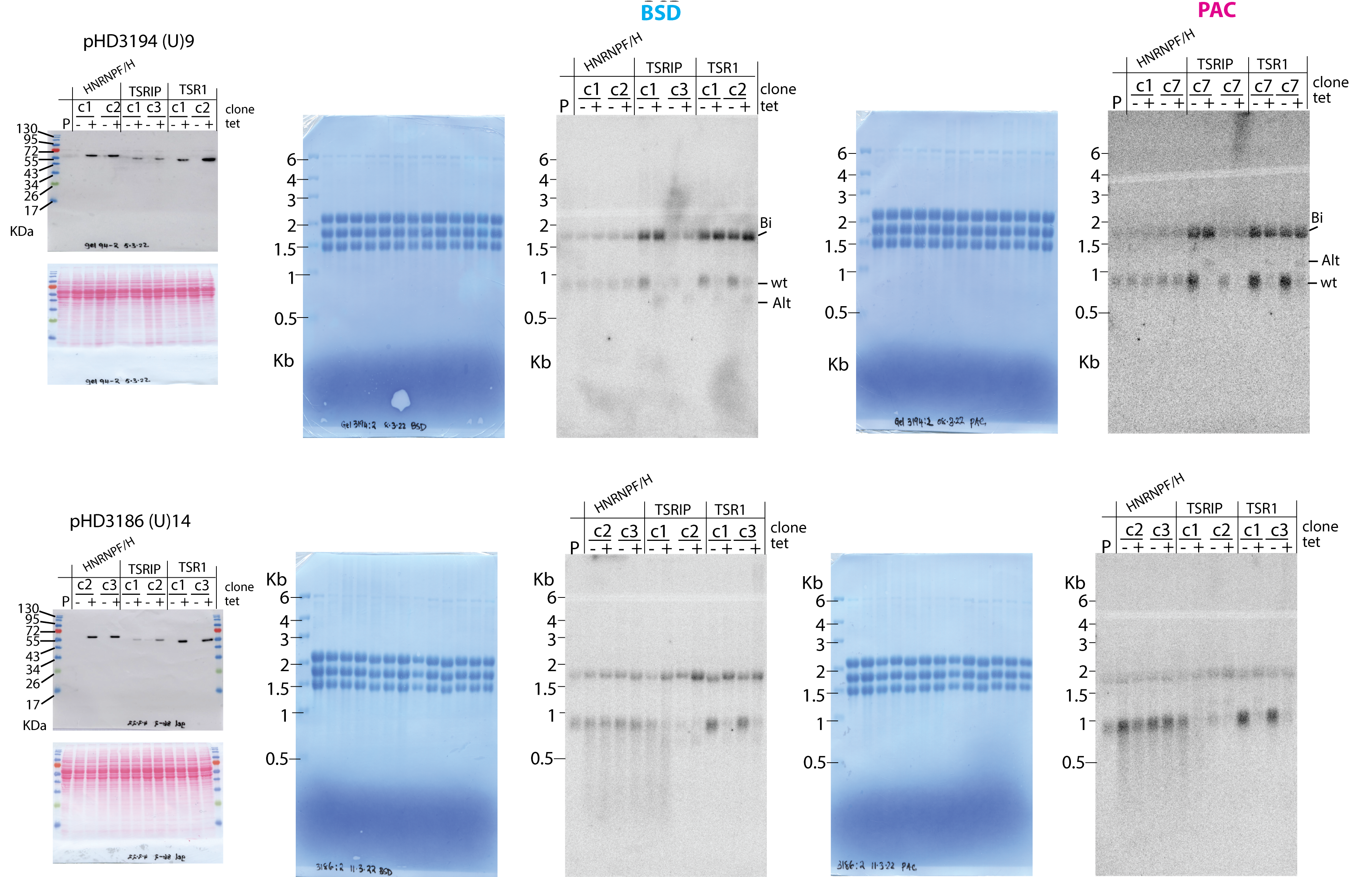

### Supplementary Figure S9

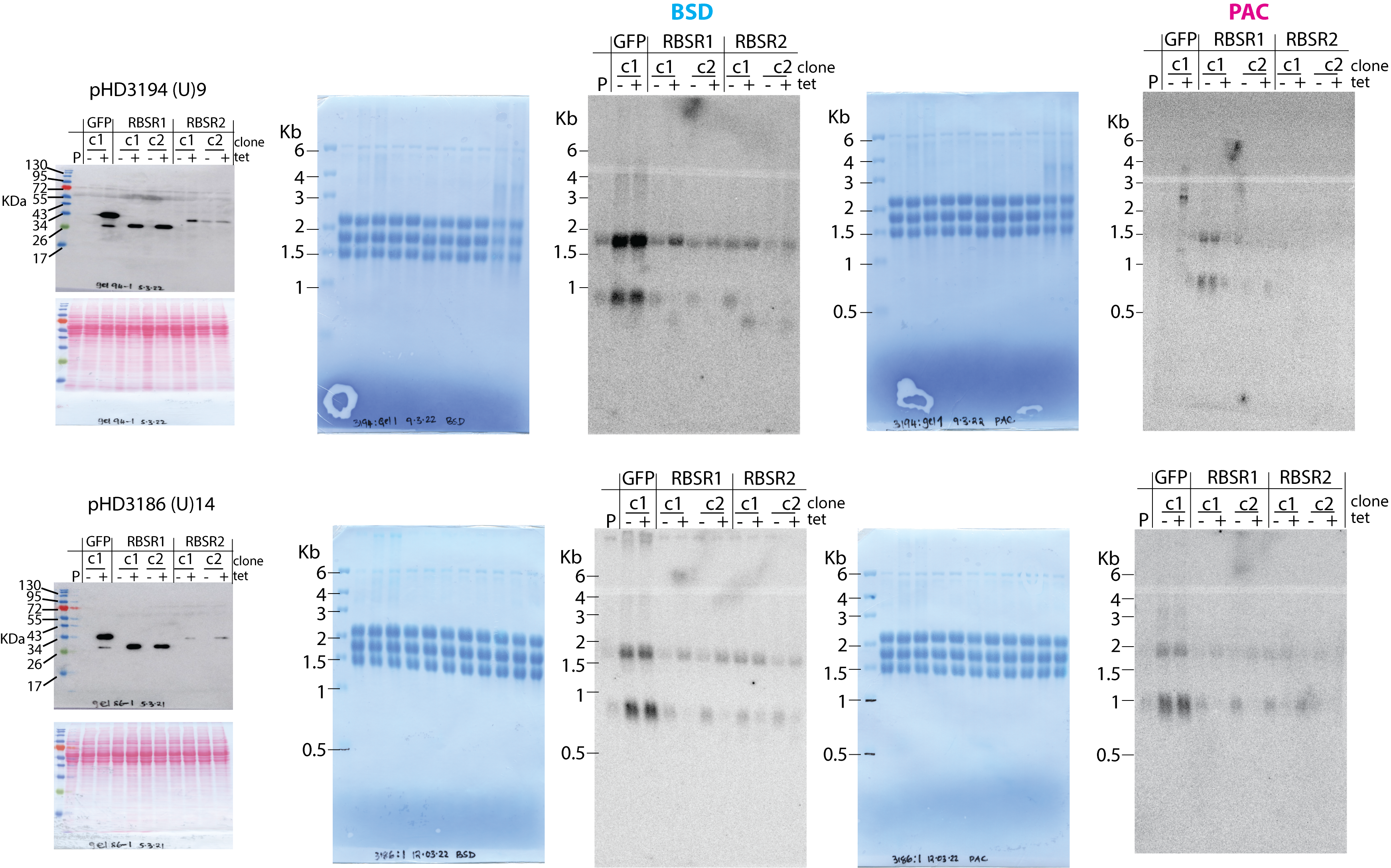

### Supplementary Figure S10

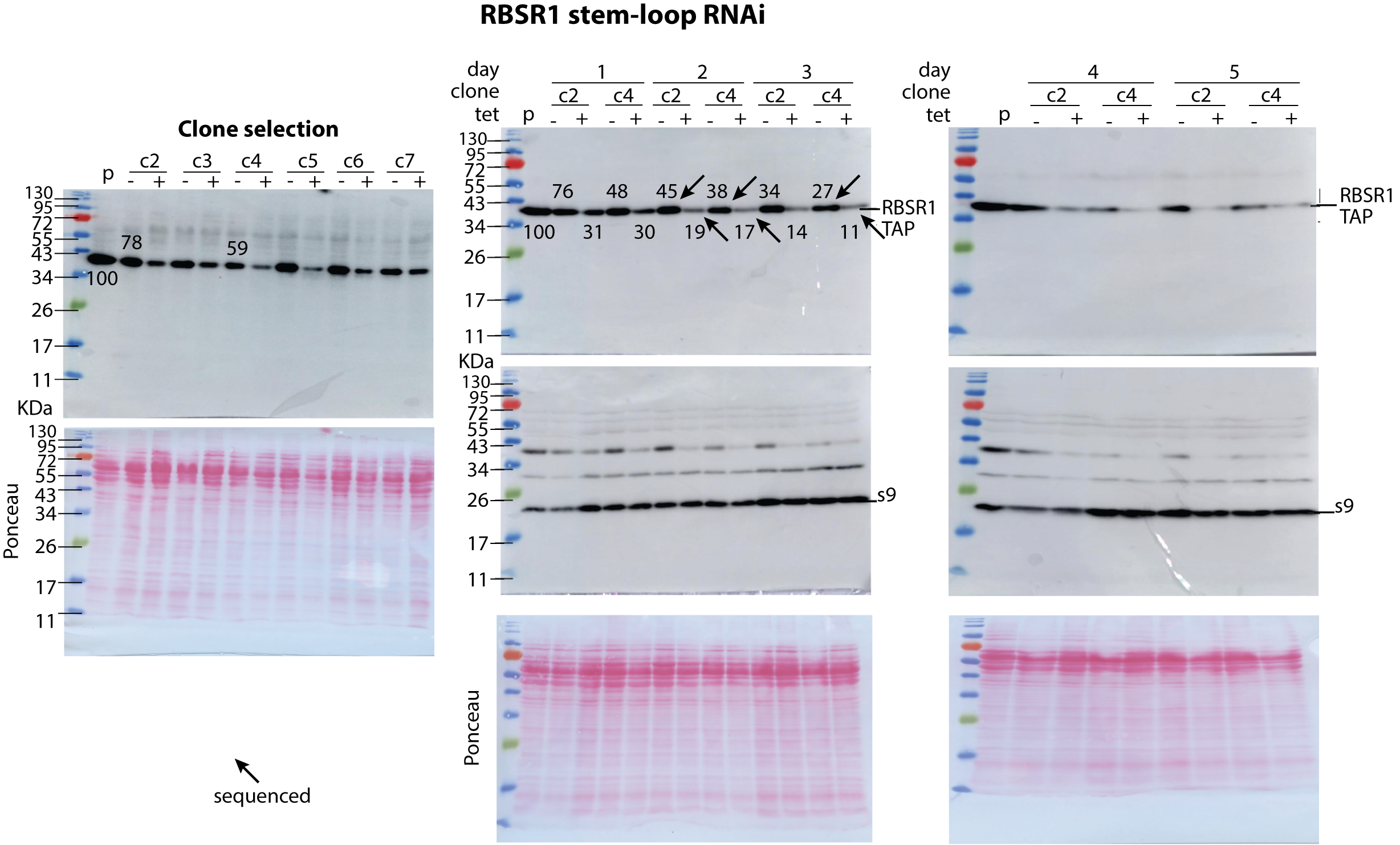

### Supplementary Figure S11

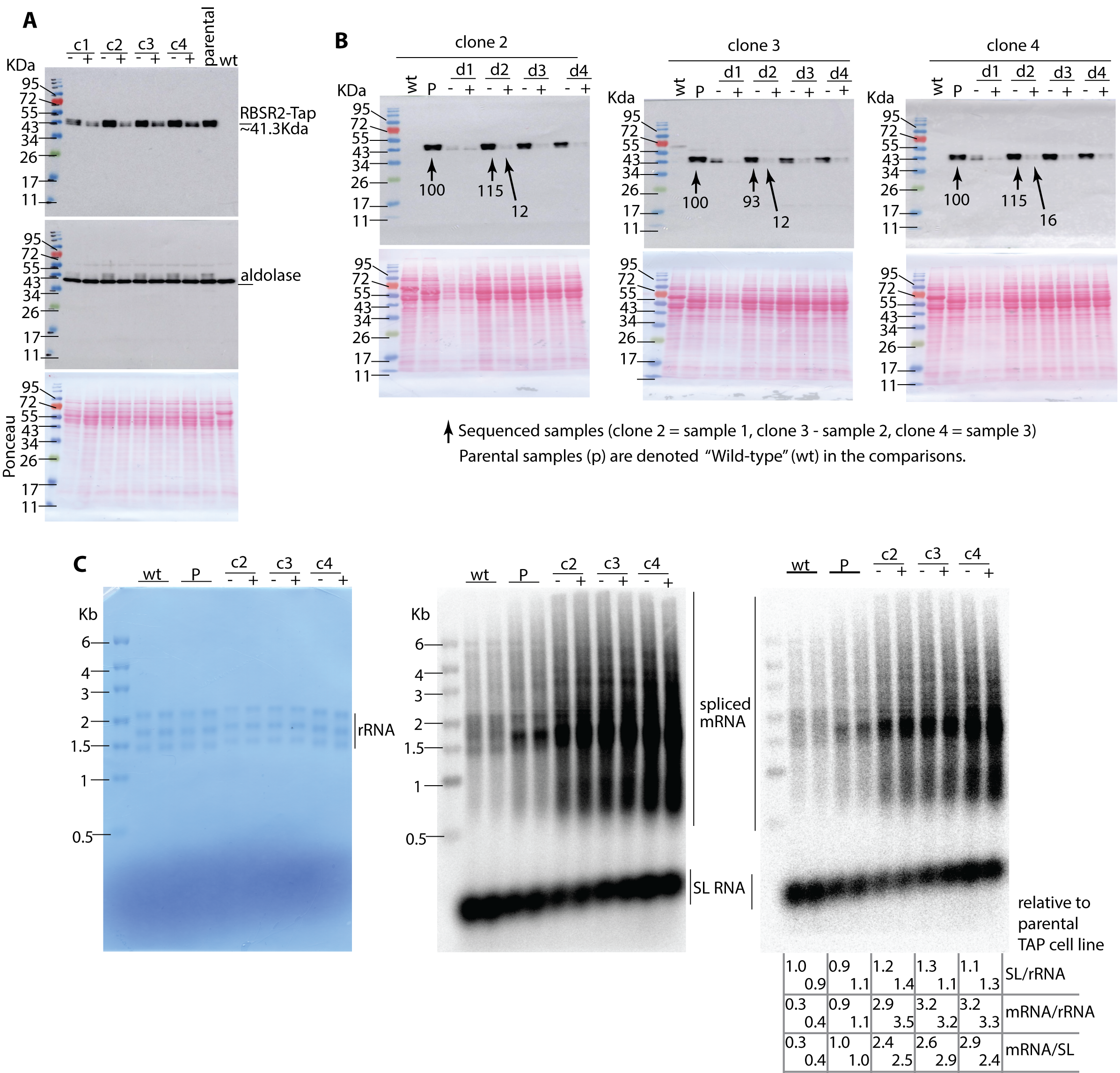

### Supplementary Figure S12

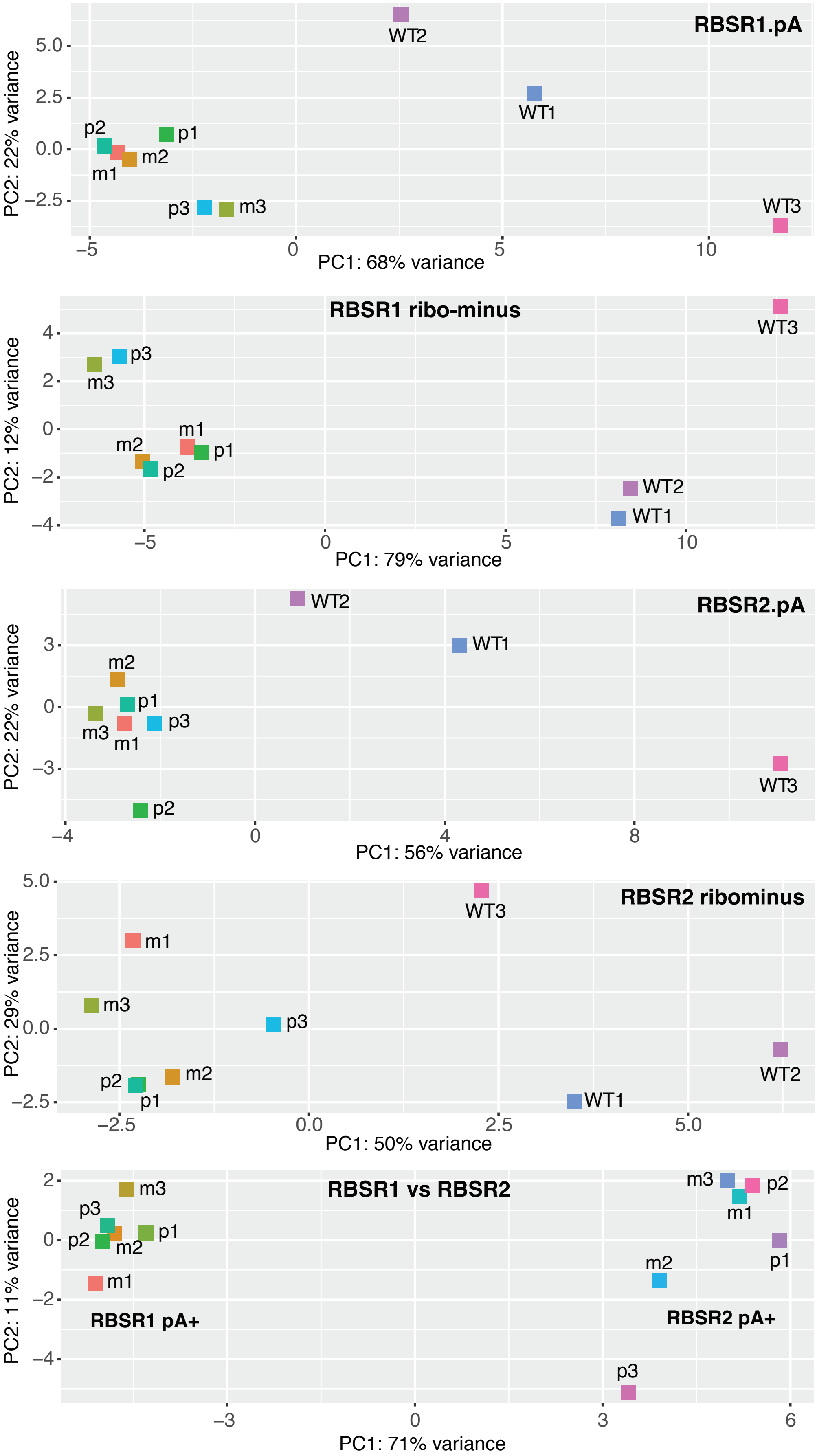


| Cell densities | % RBSR |
|---|---|
| **WT (no RNAi)** | |
| WT1: 9.3 | |
| WT2: 7.9 | (100) |
| WT3: 9.7 | |
| **RBSR1 RNAi** | |
| m1: 7.0 | 45 |
| m2: 7.5 | 38 |
| m3: 7.0 | 27 |
| p1: 12 | 19 |
| p2: 13 | 17 |
| p3: 6.1 | 11 |
| **RBSR2 RNAi** | |
| m1: 12.6 | 115 |
| m2: 9.9 | 93 |
| m3: 10.1 | 115 |
| p1: 8.2 | 12 |
| p2: 8.2 | 12 |
| p3: 8.3 | 16 |

### Supplementary Figure S13

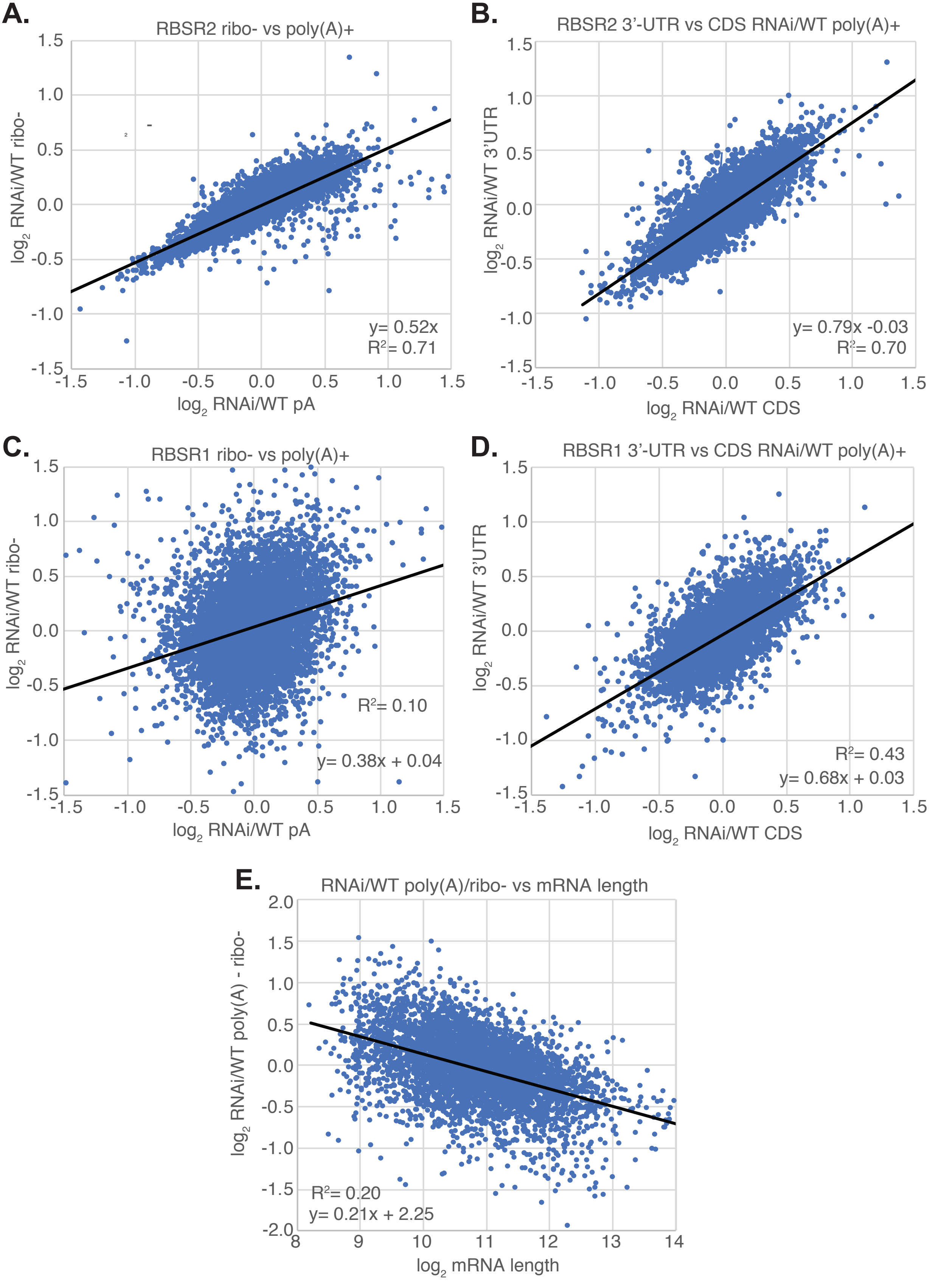
