## Supplementary Text 1 for "Sequences and proteins that influence mRNA processing in *Trypanosoma brucei*: evolutionary conservation of SR-domain and PTB protein functions"

***pHD3180 intergenic region aligned with source sequences***

**ggatccTAACACCGGGTTGTGTTGCCAAAATTGTTCTGTAGTCGCTGTGAGTTGACACGGCTAGTGCTTATGATTT**

**........ACACCGGGTTGTGTGGCCAAATTTGTTCTGTAGTTGCTGTGAGTTGACACGGCTAGTGCTTATGATTT**

**TCCTCGCGTGTGGTGCCTGTACTCAGCCCTATGCCTTATTTGCAACACATTTACGTACAGCGCACAAGAGAAGAGA**

**TCCTCGCGCGTGGTGCCTGTACTCAGCCCTATGCCTTATATGCAACACATTTACGTACAGCGCACAAGAGGAGAGA**

**AGATCACTTGAAGATAATAAATATAGGGTTGTAGGCATCTTGTTTAACTCAAATTTTCTCGTCTTGGTGTgtcgac**

**AGATCACTTGAAGATAATAAATATAGGGTTGTAGGCATCTTGTTTAACTCAAATTTTCTCGCCTTGGTGTGTCGAC**

**......GAAAATAGTATGATTGCCACCAGTTGTGTTTGATGCGTTTGTTATCTATGCAGCATTGCACAGCAAGGT**

**ATGATTGAAA.TA......GTGCCACCAGTTGTGTTTGATGCGTTTGTTATCTATGCAGCATTGCACAGCAAGGT**

**CTTCGGCACAGCAAGctcgagGCGAAAtctagaTTCATGTTTTTTTTTTTTTTACTCagatctACTTCAATTACAC**

**CTTC..................TGAAA......TTCATGTTTTTTTTTTTTTTACTCTGCATTGCAGTCT**

**...............................................................ACTTCAATTACAC**

**CACATACTACGCTTCACactagtCAGCTTACCATG**

**CA**

***Source sequences***

**>Tb927.9.8880 actinB (5'-truncated 3'-UTR)(with extra PPT T residues from Ben Amar et al, 1988).**

**ACACCGGGTTGTGTGGCCAAATTTGTTCTGTAGTTGCTGTGAGTTGACACGGCTAGTGCTTATGATTTTCCTCGCGCGTGGTGCCTGTACTCAGCCCTATGCCTTATATGCAACACATTTACGTACAGCGCACAAGAGGAGAGAAGATCACTTGAAGATAATAAATATAGGGTTGTAGGCATCTTGTTTAACTCAAATTTTCTCGCCTTGGTGTGTCGACATGATTGAAATAGTGCCACCAGTTGTGTTTGATGCGTTTGTTATCTATGCAGCATTGCACAGCAAGGTCTTCTGAAATTCATG**

**TTTTTTTTTTTTTTACTCTGCATTGCAGTCTCCGCTCTTATTTAGTTTTGCTTTACGTAAGGTCTCGTTACTGCCATAAAATA**

**EP and GPEET 3'-UTR**

**ACTTCAATTACACCAagaagtaaaattcaca**

***pHD3186 intergenic region - 3180 with 2xboxB***

**gtcgacGAAAATAGTATGATTGCCACCAGTTGTGTTTGATGCGTTTGTTATCTATGCAGCATTGCACAGCAAGGTCTTCGGCACAGCAAGctcgagCTGGGCCCTGAAGAAGGGCCCATATAGGGCCCTGAAGAAGGGCCCTATCGAAAtctagaTTCATGTTTTTTTTTTTTTTACTCagatctACTTCAATTACACCACATACTACGCTTCACactagt**

***Other intergenic regions***

**>Tb927.11.3270 Squaline monooxygenase**

**TCTAAAAATGAAGTAAAAAAGGCACGGTAAGGAGCGCCCATGGAGCTTCGAGGGCATGCCAGTTTTCATGCCCTTCTGTTTTGTGTTACAAAAGGCAAAAGTTTCTCTTTTCCTCCGTTGGCTTTCTTTTTTTTTTCCTCTGGCAGACTTTTAAGCATATAACCACTGGTACGGGAGTGAAGGAAAAAATAGAA**

**>Tb927.11.13780 profilin**

**GAGAGGAAAGAGGGTCTTCCCAATATGTGATCAAGTGTATATTACGTTATTATTATTATTATTTGCTTCTTACCGAATTTGCTACGGAGCACATGTGTATGTTTTATCGTTTCTTATTCACCTTTTCCATTAAAGAAAGAGAGCCCACAAAAAGTGAAGTTTGCCTAACTTAAGGGAAGGTTTTACTAACCGGAAATAAA**

**>Tb927.10.7420 BDF2**

**AAATATCGTTCTTGTTTCCACTTCCTTCCAGCGAAACGCTCATGATATGTTAGTTGGGCGAAAGTCTGTACATAAACATCTGTGTGTGACATGCATGTGGAAAATTTTTTCTTCGTTTCTTCACACGTGCATTTGCGTATCCACGTCTGCATAGTCGTCTATTGAGTTGAACATTGGACACGTGGGGCGGCAGCTCAAGCTCAATATATAATTTTTAGTTTTAATCTGGGAGGCGCCGGGTACCAAACAATAGCGGTCGCGGCGAAGATTCGCATACAGACGCGAGGGTGGGAAGGAGGAGGGTGAGCGAGCTCGGGCGGCGCCGAACATCTGCGTTAGAGCAA**

**>Tb927.7.940 unknown function**

**TATGGAAAAAAGTCCAAGGAAGTGGTTGAGGGAAAGAGAGTACCCTGATGCATTTATCGGTTATCGACACCCGTATGGTATGGGACGGAATCGCAGAAGTATTCCAGAGGAATCAGAGTCATCATTTTTTTTCTTTCGAAATATGATGTTTCAATGATGTCAGGAGCAGGTGAATTTCCCCACGTCACATTGTGCAGAGCATGAATTGAAAGAGAACTACTGCTCGCATATATCGGATTCTCTGCTGACTGTGCTTCTTCCTTGCTACTATTGTGTAAAATGTGACTAAGGTTAGTTTAGAAATATAATA**

**>Tb927.7.3940 MCP 16**

**CAAACATATGTGTTTGAATACATATGCGTGTGTGCTTGAAGAATCAGGTTTCTCAACCTTGTTGAGCCCGGAACGAGCACCTATCTTTGCTTCCTCTTCCTTGTTTTGGGTGGAACTCCCCCACCCTTTGGATGCTCGTTCGTCCGTTCGGTTCCACTATTTTTTTTCTTTGCATGTTTGTGCTACCGAGGGTCGAAGGGGTAAGATAGGAAAATCTACCTGTTAGGACGGTGCCTGCGGAATATACGTGTGACAGAGAAAAGTAGCGTTGGGTGCGAACACCGAGAGAGTATCTGTCATCAATTCAGATCTCGGCGAGTAACATTTATCGAATAAGACCTGTCGTACTCGTCACTCACTCTGTGGCAGAATACAAATG**
