## Supplementary for "Sequences and proteins that influence mRNA processing in *Trypanosoma brucei*: evolutionary conservation of SR-domain and PTB protein functions"

**Supplementary material**

**FIGURES**

*Figure S1*

Raw data for Figure 5 and Figure 6: Western blot, full Northern blots with replicates for two clones. and puromycin susceptibility curves labelled according to clone, for tethering of GFP and HNRPH/F

*Figure S2*

Raw data for Figure 5: Western blot, full Northern blots with replicates for two clones. and puromycin susceptibility curves labelled according to clone, for tethering of CPSF3 and SF1.

*Figure S3*

Raw data for Figure 5: Western blot, full Northern blots with replicates for two clones. and puromycin susceptibility curves labelled according to clone, for tethering of CPSF65.

*Figure S4*

Raw data for Figure 6: Western blot, full Northern blots with replicates for two clones. and puromycin susceptibility curves labelled according to clone, for tethering of DRBD3 and DRBD4

*Figure S5*

Western blot, full Northern blots with replicates for two clones. and puromycin susceptibility curves labelled according to clone, for tethering of TSR1 and its interaction partner TSRIP.

*Figure S6*

Western blot, full Northern blots with replicates for two clones, and puromycin susceptibility curves labelled according to clone, for tethering of RBSR1 and RBSR2.

*Figure S7*

Western blot, full Northern blots for *BSD* and *PAC*, with replicates for two clones, for tethering of DRBD3 and DRBD4 to the (U)_9_ and (U)_14_ boxB reporters.

*Figure S8*

Western blot, full Northern blots for *BSD* and *PAC*, with replicates for two clones, for tethering of HNRNPF/H, TSR1 and TSRIP to the (U)_9_ and (U)_14_ boxB reporters.

*Figure S9*

Western blot, full Northern blots for *BSD* and *PAC*, with replicates for two clones, for tethering of RBSR1 and RBSR2 to the (U)_9_ and (U)_14_ boxB reporters.

*Figure S10*

Western blots showing levels of RBSR1-TAP with and without RNAi induction. "P" is the precursor cell line with the TAP tag but no RNAi plasmid. The numbers are quantitation and the arrows indicate samples used for RNASeq.

*Figure S11*

Western blots showing levels of RBSR2-TAP with and without RNAi induction are in A and B. "P" is the precursor cell line with the TAP tag but no RNAi plasmid. The numbers are quantitation and the arrows indicate samples used for RNASeq. S9 is ribosomal protein S9. Panel C shows a Northern blot of the RNA used for seequencing, hybridised with a spliced leader probe. "P? is two samples of the input (tagged) line that serves as the "wild-type" control in the RNASeq analysis. "wt" is RNA from another experiment; the low amount of mRNA in these might be caused by high cell density but this is uncertain. Two exposures are shown and relative quantitation of the shorter exposure (normalised to rRNA) is shown.

*Figure S12*

Principal component analysis for RNASeq of the RBSR1 and RBSR2 RNAi lines. "WT" here refers to the line expressing RBSR2-TAP, without any RNAi plasmid. "m" means minus tetracycline, "p" means plus tetracycline, for the three replicates inlustrated in Figures S10 and S11. Cel densities (multiplied by 10^-5^) and the percent of the RBSR protein for each sample (as shown in Figures S10 and S11) are also shown. "pA" is poly(A)+ RNA, ribominus is rRNA depleted.

*Figure S13*

Scatter plots comparing the RNASeq datasets. In all cases the results for +tet and -tet were pooled. All results are log2-transformed.

A. RBSR2, RNAi cell line/WT, poly(A)+, on x-axis, ribo-minus on y-axis.

B. RBSR2, RNAi cell line/WT, poly(A)+, coding sequence (CDS) on x-axis, 3'-UTR on y-axis.

C. RBSR1, RNAi cell line/WT, poly(A)+, on x-axis, ribo-minus on y-axis.

D. RBSR1, RNAi cell line/WT, poly(A)+, coding sequence (CDS) on x-axis, 3'-UTR on y-axis.

E. RBSR1, RNAi cell line/WT, poly(A)+ result divided by ribo-minus result R on y-axis, mRNA length on x-axis.

**TABLES**

*Table S1*

Plasmids and oligonucleotides.

*Table S2*

PPT screen. For details see first sheet.

*Table S3*

Mass spectrometry data. For details see first sheet.

*Table S4*

Abundances of proteins implicated in splicing, downloaded from Tinti, M. and Ferguson, M. (2022) *Wellcome Open Research*, **7**.

*Table S5*

Transcriptome results: raw reads.

*Table S5*

Transcriptome results: DeSeq2 analysis.

**DNA Sequences**

*Supplementary text 1*

Intergenic sequences used for Figure 3.

*Supplementary file 1*

pHD3180 sequence, ApE format.

*Supplementary file 2*

pHD3186 sequence, ApE format.

*Supplementary file 3*

pHD3190 sequence, ApE format.

*Supplementary file 4*

pHD3259 sequence, ApE format.

*Supplementary file 5*

The "Wild-type" *BSD* mRNA sequence, as DNA in ApE format.

*Supplementary file 5*

The "Wild-type" *PAC* mRNA sequence, as DNA in ApE format.
